# Effective elasticity and persistence of strain in active filament-motor assemblies

**DOI:** 10.1101/2021.12.14.472714

**Authors:** Arvind Gopinath, Raghunath Chelakkot, L Mahadevan

## Abstract

Cross-linked composite filaments animated by molecular motors are ubiquitous in biology. An important length scale that controls the range of force transmission in these filament-motor assemblies is the distance over which elastic strain fields persist. Previous studies have identified this persistence or decay length in passive networks, and recently, in random elastic networks with embedded elements generating constant forces or stresses. Yet, the range over which strains persist in assembles comprised of motors attaching stochastically to nearly inextensible filaments remains unclear. Here, we analyze the intrinsic decay length for extensional strain in a minimal model – ordered composite assemblies comprised of cross-linked filaments interacting with arrayed motors. We explores three related concepts: the effective active elasticity of these assembles, the length scale over which localized elastic strains persist in these systems, and its dependence on motor kinetics for static and oscillatory deformations. By combining mean-field theory for noiseless weakly extensible systems with Multi-Particle Collision Brownian simulations for noisy systems we are able to additionally investigate the effects of motor noise. For steady strains, we find that the persistence length is independent of motor kinetics and dependent on the passive elasticity of the composite. For oscillatory strains, the persistence length is frequency dependent and depends on passive elasticity and on motor kinetics. For strong motor activity, the decay length is significantly modulated by extensional softening resulting from attached motors. Theoretical predictions agree qualitatively with our simulations, even at low motor numbers or moderately strong motor noise. Taken together, our results highlight how extensional strain and correlated dynamics in possess a finite range of influence even in active networks comprised of nearly inextensible filaments.

## 1 Introduction

### 1.1 Motivation

Active strain generated by molecular motors that interact with biofilaments and networks is the main mechanism of force generation and motion in cell biology ^1–11^. In naturally occurring filament-motor assemblies and networks, and in reconstituted versions of these, filaments are dynamically stretched or bent by forces imposed by motors. The strength of the deformation depends on the intrinsic elasticity of the filament, geometric constraints, intrinsic motor kinetics, active elasticity imparted by attached motors, and noise. Noise can have either thermal or athermal origins. Thermal noise is a consequence of fluctuations caused by the ambient medium. Athermal or motor noise is a consequence of fluctuations in the number of attached motors. Since temporally varying filament deformations can in turn regulate motor activity ^12–15^, the dynamics of the overall filament-motor assembly is a consequence of the two-way nonlinear coupling between motor mechanochemistry and filament elasticity. Ultimately, it is the spatial extent of correlated activity in these systems that both enables and limits function and physiology. This length-scale which one can interpret as the persistence or decay length, quantifies the range over which locally generated or imposed deformations remain correlated.

Previous studies have identified the persistence length in passive networks, and have provided clarity on the role of the shear modulus in controlling this length. Because many biofilaments, such as microtubules, are intrinsically stiff in the absence of cross-linking filaments, filament extensibility is usually neglected in studies of cross-linked filamentous networks. However, recent models of passive filament bundles ^7,16–18^ account for finite stretching stiffness of the filaments. Experiments indicate that when such bundled filaments are bent, the length over which mechanical information is transmitted is determined by a combination of the effective shear stiffness of the bundle and the stretching and bending stiffness of individual filaments. In particular, work by Ward *et al*. ^18^ on mutually sliding actin filaments suggests that the non-trivial dependence of sliding friction on the filaments’ overlap length can be explained by a length scale arising from the finite extensional rigidity of f-actin. More recent studies have focused on stress propagation in soft media such as active gels ^19–21^, active polymers ^22–26^, and disordered filament networks. Interestingly, actuation of selected modes rather than simple screened decay of displacement fields has been predicted in a class of active solids ^27^. Persistence lengths are also crucial in the context of cell motility, multicellular assembly, and durotaxis. For instance, non-adherent cells, such as crawling fibroblasts exert localized dipolar stresses on the substrate, and are hypothesized to mechanically communicate with neighboring cells through induced strain fields with finite range ^28–30^.

In summary, previous work has primarily focused on identifying the persistence length in passive networks, or in random elastic networks with embedded elements generating constant forces or stresses. However, the emergence of analogous decay lengths arising in assemblies comprised of nearly inextensible filaments interacting with motors generating state- or time-dependent forces remains unclear.

Consider Figure 1 that depicts a composite filament-motor assembly subject to active forces from two distinct motor aggregates (labeled I and II). If the filament is truly inextensible, mechanical activity by group (I) and shear (sliding) induced by this patch is transmitted by attached motors in (II) over effectively infinite inter-aggregate distances. The crucial idea here is that any manner in which softness is introduced into system elasticity provides a means by which this unphysical scenario is prevented. Indeed it is clear that when the filament is very soft (extensible), the ability of motors in group (I) to mechanically communicate with group (II) via the intervening elastic filament is strongly limited.

**Fig. 1.**
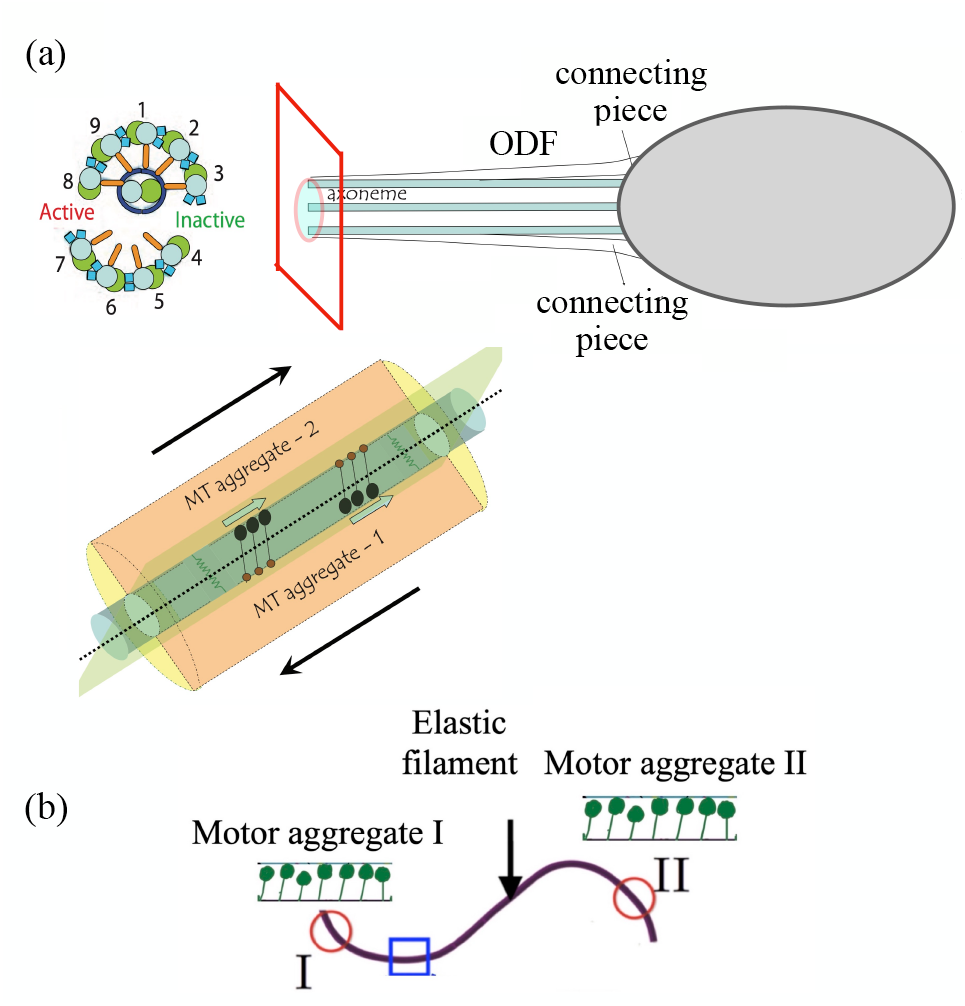
(a) A schematic of the cross-section of the slender axoneme – a canonical structured motor-filament assembly. The connecting piece connects the head with the membrane enclosed (9+2) doublet-microtubule structure that runs along the length of the axoneme. Interacting with the 9 doublet pairs (left) are axonemal dynein molecular motors organized into inner and outer dynein arrays. Each array comprises of motors that attach stochastically to adjacent microtubule doublets, form temporary cross-bridge structures, generate forces and cause relative sliding. Studies suggest that during the beating cycle, the axoneme assembly may organize into two semi-cylindrical sections sliding past each other due to the action of dynein arrays motivating the minimal geometry studied here. (b) Schematic of a motor-filament aggregate with spatially separated motor groups (I) and (II) interacting with an elastic filament backbone(a segment of which is highlighted in blue). The softer the backbone, the lower the ability of the motor groups to sense each other.

This observation is particularly pertinent to the oscillatory beating of eukaryotic flagella and cilia. The eukaryotic flagellum is a highly ordered, composite organelle composed of stiff filaments (microtubules), active and elastic motors (axonemal dyneins), and soft passive elastic linkers (nexins) ^13–15^ enclosed in a sheath. The structural and functional aspects of the axonemal components and the modes of activation to induce beating have been extensively investigated ^31–40^. Models with varying degrees of complexity have been introduced to study the dynamics of and deformations in oscillating flagella; these include curvature control models, shear control models and buckling-induced activation controlled models ^8–15,41–56^. In all these theories however the axoneme is assumed to be inextensible. This is usually justified by the very high extensional modulus of the doublet microtubules comprising the backbone of the axoneme ^3,51^. Consequently, axial strain and extensional deformation along the axoneme are neglected. The assumption of inextensibility is certainly reasonable for short cilia, as in the green alga *Chlamydomonas reinhardtii* with cilia of length ~ 10 *μ*m). Theories assuming an inextensible axoneme predict that beating wavelength scales with the length of the flagella; thus very long flagella will have correspondingly long wavelengths. This contrasts strongly with experimental evidence that suggests that flagella beating wavelengths are self-limiting ^2–4,51^. In long flagella ^51^ such as rat sperm flagella (lengths ~ 170-180 *μ*m), quail sperm (lengths ~ 230-250 *μ*m) and *Drosophila* sperm (lengths ~ 1000 *μ*m or longer), substantial degradation of mechanical information may be expected as one moves from the proximal end to the distal end.

Taken together, experimental evidence therefore strongly suggests that the finite extensibility of the microtubules, the mechanical state of the flagellum, and the motor-mediated elasticity of the assembly itself must be considered to understand flagellar dynamics ^51^, and other similar filament-motor assembles. Here, we study this further in a minimal system that allows us to identify and dissect individual mechanisms impacting the persistence and decay of elastic strain. Specifically, we address the following questions:

1. *What is the effective coarse-grained elasticity of composite arrayed active filament-motor assembles?*,
2. *What sets the length scale over which localized elastic strains persist in such systems?, and*
3. *How does this persistence vary with motor kinetics in both static and oscillatory settings?*

Referring back to Figure 1 and recognizing that motor groups interacting with an assumed infinitely stiff filament will eventually coordinate, we hypothesize that elasticity, be it intrinsic (passive) extensional elasticity or active elasticity and associated shear softening, can mediate interactions between spatially separated regions of the filament. Additionally, kinetics of motor activity set by ATP hydrolysis rates can also be coupled to effective extensibility by introducing shear stiffening or extensional softening.

The paper is organized as follows. In Section 2, we elucidate the coupling between bending, shearing, and extensional elastic components in a passive composite containing filaments and passive cross-linkers. In Section 3, we use these results as a basis to formulate and analyze a mean-field model for the active filament-motor composite when the influence of noise is negligible and the motor density is large. We obtain a coarse-grained description of the elasticity of these assemblies and identify passive and motor-driven active stochastic contributions to the overall elastic properties. Using this model, we then study how localized and oscillatory extensional and shear strains decay from the point where they are imposed and present analytical expressions for the persistence length for each case. For steady strains, we find that the persistence length is independent of motor kinetics but dependent on the passive elasticity of the composite. For oscillatory strains, our analysis highlights the crucial role played by the ratio of the imposed frequency to the detachment rate in determining and modulating the persistence length. In Section 4, we analyze a discrete stochastic numerical model that incorporates motor noise arising due to discrete attachment and detachment kinetics and the finite number density of motors. Theoretical predictions agree qualitatively with our simulations, even at low motor numbers or moderately strong motor noise. Taken together, our results highlight how extensional strain and correlated dynamics in possess a finite range of influence even in active networks comprised of nearly inextensible filaments.

## 2 Persistence lengths in passive assemblies

We start our analysis by studying a passive composite filament comprised of two strips that are connected by permanent elastic cross-links. The elastic properties of such a passive composite assembly can be quantified by its bending, shear, and extensional moduli, which are mutually coupled via the geometry of the composite.

In Figs. 2(a) and 2(b) we illustrate our minimal model: a composite filament composed of two cross-linked elastic strips (top strip in green). Both strips have a rest length *ℓ*, a thickness *w* ≪ *ℓ* and a lateral width *b* ≫ *w*, and are maintained a distance *D* ≪ *ℓ* apart by a series of regularly spaced passive linear elastic cross-links (springs) with areal density *ρ*_N_ and spring stiffness *k*_N_. These springs, while compliant to shearing forces, are inextensible and thus prevent any change in the thickness of the gap *D*. We focus on planar bending deformations, that is, one-dimensional sliding and stretching deformations of this composite filament. The filament+motor system is henceforth simply referred to as an *assembly*. There are no deformations along the neutral axis, and therefore the deformation fields are solely functions of the arc-length parameter *s* ∈ (0, *ℓ*) specifying the location of the material points on the strip along the axial direction. At length scales larger than the spacing between passive springs, the assembly acts as a composite filament of thickness *D*, with an effective shear modulus that depends on *D, ρ*_*N*_, and *k*_*N*_.

**Fig. 2.**
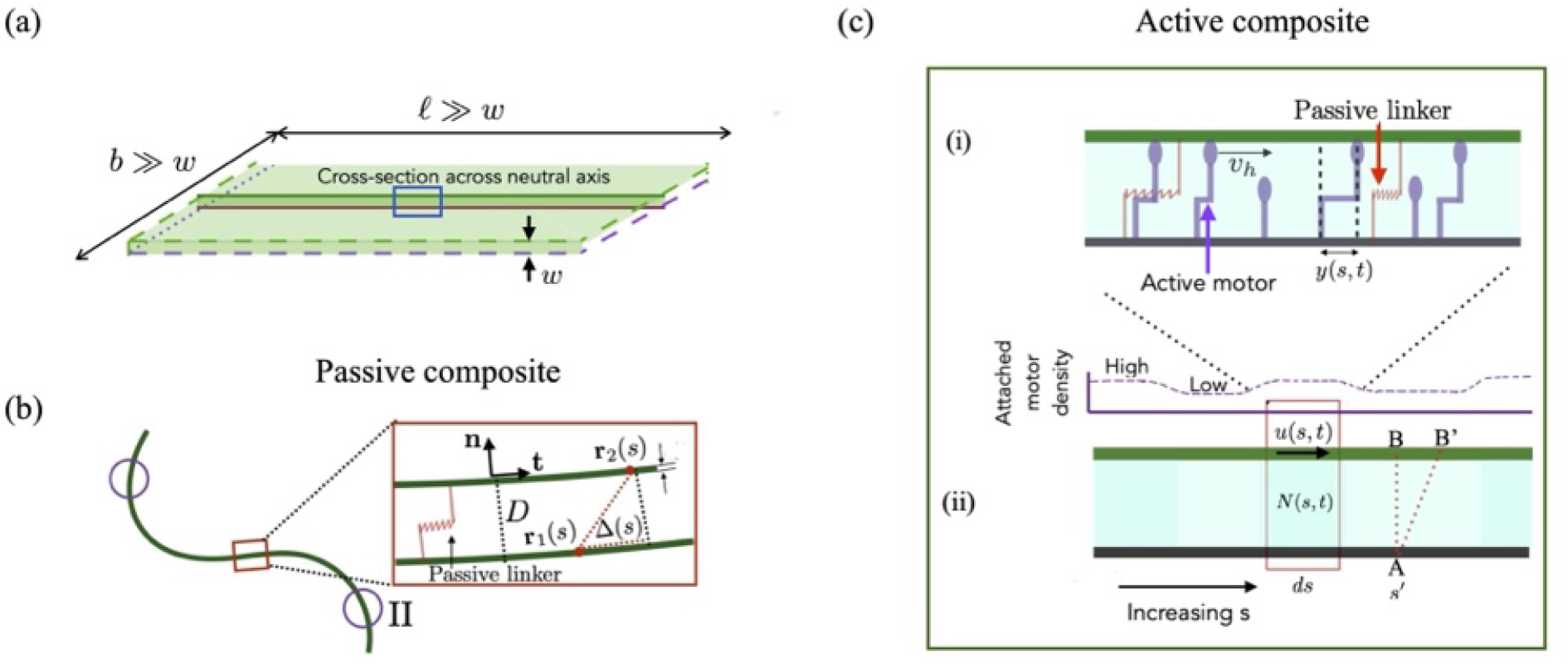
(a) Schematic of the composite strip with extensional and shear deformations analyzed in the mean-field model. We consider two strips (surfaces) each of length *ℓ*, thickness *w* ≪ *ℓ*, and lateral extent *b* ≫ *w*, and separated by a distance *D*. We choose *b* ≪ *ℓ*, in this limit the transverse direction may be treated as a neutral direction and one may just study 1D deformations in the direction along the long axis (lengthwise direction). A small portion of the composite filament in this direction is marked in blue and is magnified in cases (b) and (c). (b) Passive composite system comprised of two stretchable sheets or filaments connected by passive crosslinks. Bending is restricted to a single preferred plane as shown. The cross-links maintain the distance *D* between the two filaments and provide shear resistance. (c) Active motor-filament composite with molecular motors. (i) (Microscale) The bottom surface/filament (purple dashed lines) is rigid and fixed. The upper surface/filament (green dashed line) can stretch and slide, but cannot bend. Also shown are permanently attached passive elastic links (red), transiently attached active motors (purple). Motors attach with a pre-extension *d*_*m*_ and move along the upper filament with constant speed *v*_*o*_ relative to their base. The mechanical stressed state of attached motors is quantified by their internal extension *y*. (ii) (Macroscale) Coarse-graining the motor activity, we derive a continuum description for the local density of attached motors *N*(*s, t*) and the local mean motor extension *Y* (*s, t*). Shown is a typical deformation that is a combination of slide/shear and extension and moves material point *B* to *B*^′^.

Consider now a shear stress *σ*_*s*_ that acts on the upper filament on an effective surface of area *bℓ*, and causes a sliding displacement Δ between the filaments. This applied stress is supported by the (elastic) stress exerted by the connecting springs *ρ*_N_*bℓ*. The balance of forces then yields the scaling relationship *σ*_*s*_*bℓ* ~ Δ*k*_N_*ρ*_N_*bℓ* ~ *G*Δ*bℓ*. The effective shear modulus of the entire strip that includes contributions from these passive springs is *G*^∗^ ~ *Dk*_N_*ρ*_N_. For a strip of lateral width *w* corresponding to the neutral direction, we estimate an effective stiffness *G*^∗^*w/D* ~ *wk*_N_*ρ*_N_ consistent with previous analyses of a passive cross-linked railway track model.

We note that this analysis, and the theories developed in sections §2 and §3, are applicable to assemblies comprised of cross-linked planar strips or cross-linked semi-cylinders (as shown in Figure 1(b)), after appropriate rescaling of elastic properties.

### 2.1 Energy functional incorporating all deformation modes

Consider two points **r**_1_ and **r**_2_ as shown in Fig. 2(b) that face each other when the filaments are in the undeformed state. As the composite filament deforms, the location of **r**_2_ relative to **r**_1_ changes. The slide Δ(*s*) between the points is defined as the difference between **r**_1_ and **r**_2_, relative to the initial value before deformation. The angle made by the local tangent at *s* to the centerline, *θ* (*s*), is related to Δ(*s*) and the displacement field, *u*(*s*), by geometry. The local shape is quantified by the relations

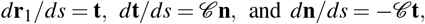

where **n** and **t** are the normal and tangent vectors evaluated at **r**_1_, and *C* = *d***t***/ds* · **n** ≈ *dθ/ds* is the local curvature. Using (*D* + *w*)*/ℓ* ≪ 1, we write to leading order

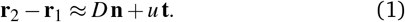

We now consider deformations with the tangent angle at *s* = 0 set to zero. Differentiating (1) with respect to *s* yields

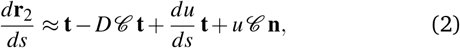

For small deformations, *u C* ≈ 0 in (2). Since

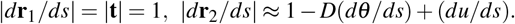

Stipulating

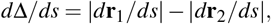

we obtain a simplified version of (2) that relates the spatial variation in the shear displacement to spatial variations in bending and stretching,

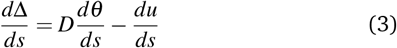

Integrating (3) from *s* = 0 to *s* using the boundary condition *θ* (0)= 0 produces the relationship between shear, extensional, and bending deformations,

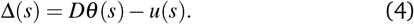

Taking into account the all three modes of deformation, we write the total elastic energy functional *E*_*T*_,

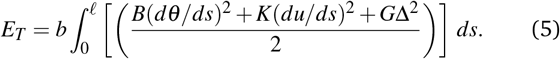

Here, *B* ~ *Ew*^3^, *K* ~ *Ew* and *G* ~ *ρ*_N_*k*_N_ are the passive bending, stretching, and shear moduli, with *E* being the Young’s modulus.

### 2.2 Minimizing the energy functional yields persistence lengths

The Euler-Lagrange equations obtained by minimizing (5) lead to force-balance relations that provide static solutions of the state and shape of the deformed filament, that is, the fields Δ(*s*) and *θ* (*s*), which we derive next.

As bending stiffness *B* → ∞, *dθ/ds* → 0, and the filament deforms such that Δ ≈ −*u*. The minimization of (5) then yields

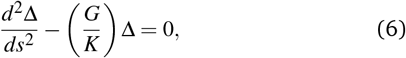

from which we deduce the relaxation length scale for pure extension

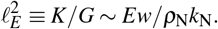

Next, we consider the limit of inextensible filaments, K → ∞ and *du/ds* → 0. Using the relation Δ ≈ *Dθ*, and minimizing (5) we get

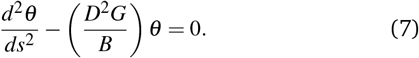

Thus, the persistence scale for pure bending is given by 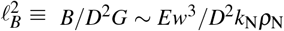. Physically, this implies that as the density of passive links or their stiffness increases, the persistence length associated with purely bending modes decreases.

Finally, we consider the general case where both the bending and the stretching modes are allowed. Allowing for sliding deformations at *s* = 0, we retain the bending, extension, and sliding terms in (5), and repeat the minimization setting *δ E*_*T*_ */δθ* = 0 and *δ E*_*T*_ */δ* Δ = 0 to obtain the coupled equations;

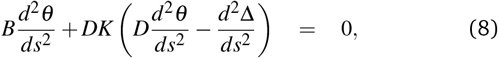

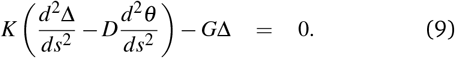

Eliminating *θ* from this pair, we get

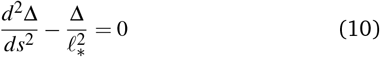

with the persistence length *ℓ*_∗_ given by

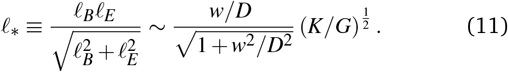

Equation (11) demonstrates that, for fixed geometric factors *w* and *D*, the persistence length 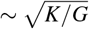 depends on the independently tunable parameters *K* and *G*.

An important feature here is the dependence of the shear modulus *G* on the density (number) of crosslinkers. Shear stiffening as a result of high linker density effectively softens the composite filament. The generality of the framework suggests that similar length scales should appear in a variety of soft systems where bending, shear, and extension are coupled, as in fact has been observed more than 75 years ago in both microscopic and macroscopic contexts ^57,58^.

### 2.3 Estimates of the persistence length in inactivated cilia

To put the discussion in perspective, we consider typical parameters for a typical eukaryotic axoneme (ciliary axoneme). We set *w* to scale approximately as the radius of a microtubule doublet (*w* ~ 20 nm), *D* to scale as the spacing between the doublets (*D* ~ 40 nm ^12^), *E* to roughly scale as the Young’s modulus of microtubules (*E* ~ 1.2 GPa ^3,12^), and *k*_N_ to scale as the stiffness of passive nexin links ^53^ (*k*_N_ ~ 16 − 100 pN *μ*m^−1^). The density of passive links *ρ*_N_ is evaluated using ^53,54^ *wρ*_N_ ~ 10^5^ − 10^7^ m^−1^.

Ignoring the active motors and retaining only the effect of the permanent nexin cross-links, we estimate *ℓ*_*E*_ ~ 200 − 400 *μ*m with *ℓ*_∗_ ~ 80 − 200 *μ*m. Of course, in beating flagella and cilia, motor activity dominates, and transiently attached motors will contribute to the overall shear resistance and modify this preliminary estimate. An upper bound that incorporates the effects of attached motors is obtained by assuming that all motors are attached. Using estimates for motor density *O*(10^8^) m^−1^, and effective motor stiffness ~ 10^−3^ N/m, we estimate persistence lengths *ℓ*_*E*_ ~ 5 − 10 *μ*m highlighting how shear stiffening by active motors makes the composite softer and decreases the length scales over which extensional and shear deformations decay. Cilia are typically around 5 − 15 *μ*m in length, while the flagella of the eukaryotic spermatozoa are often longer than this scale ^3^, suggesting that filament elasticity modulates persistence lengths. We will re-visit these estimates for the persistence length after the analysis presented in sections §3.

## 3 Mean-field theory for active assemblies

Having analyzed the decay length in a model passive composite assembly, we now move on to the case of active filament-motor assemblies. The composite we analyze here in the mean-field model is illustrated in Figs. 2(a) and 2(c). The system consists of two cross-linked filaments (denoting flat strips as shown in Fig. 2(a), or semi-cylinders as in Fig. 1(b)) with unit width perpendicular to the plane of the figure. These filaments are held a distance *D* apart by passive springs that are compliant in shear. To further simplify our analysis, only the upper filament is considered extensible; the lower filament is straight and inextensible. We will henceforth focus on the competition between shear and extension in setting decay lengths, and bending modes are henceforth ignored. Finally, motivated by the ubiquitous slender structures in biology, we restrict ourselves to cases where *b* ≫ max(*w, D*). As before, extensional and shear deformations (strains) fields are functions of the axial arc-length parameter *s*.

In addition to passive springs, the gap between the upper and lower filaments is populated by arrayed molecular motors that are grafted to the lower filament and distributed uniformly (homogeneously) along *s* with constant density *ρ*_*m*_. The action of these motors is to generate an active force *F*_*m*_(*s*) (per unit width) that displaces the upper filament relative to the lower filament. Microscopically, this force arises as motors periodically attach and exert tangential forces on the upper surface. The upper filament, lower filament, passive springs, and molecular motors together constitute the filament-motor aggregate.

### 3.1 Global force balance

Consistent with previous studies, we specify that each attached active motor behaves elastically as an active linear spring ^9,10^ with spring constant *k*_*m*_ and zero rest length. The internal mechanical state of the motor is characterized by a single variable – its intrinsic extension *y*(*s, t*) – that quantifies the displacement of the motor head that is attached to the upper filament relative to its base attached to the lower filament.

To make progress towards a mean-field theory, we ignore Brownian diffusion of the motors and further specify a simple model for the attachment and detachment process. The motors attach to the filament in a pre-strained state with extension *d*_*m*_. Once attached, to relax the pre-stress, the head is dragged by the moving filament and also slides relative to it, resulting in a changing motor extension *y*(*s, t*). This sliding speed depends on the strain (extension) experienced by the motor. Attached motors undergo a conformational change and, when attached, exhibit a motor strain (extension) given by

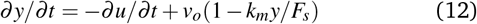

where *v*_*o*_ is the maximum velocity (at zero load) of the motor and *F*_*s*_ is the stall force at which the motor velocity vanishes. Although a non-linear force-speed relationship has also been proposed in the literature, here we study the linear form as suggested by Eqn. (12). An attached motor exerts a tangential force on the filament, here ∝ *k*_*m*_*y*. This interaction force between the motor and the filament also results in motor detachment by affecting the detachment probability. We also note the two-way coupling between motor extension and the continuum displacement field *u*(*s, t*) in Equation (12).

Let the total density of motors *ρ*_*m*_ be large enough to allow a mean-field description of the motor states and the extension of the filament. At these high densities, the number of attached motors is large enough that fluctuations in the time average of the density motors *ρ*_*a*_(*s, t*)= *ρ*_*m*_*N*(*s, t*) are small compared to the mean value. In this limit, motor extension (stretch) *y* can be treated as a continuous variable and can be used to define ensemble averages. Specifically, we can formally define the average of the variable *X* via ∫ *XP*_*a*_*dy* ≡ ⟨ *X* ⟩ *Nρ*_*m*_. Consider now an infinitesimal length *ds* of the filament centered around location *s*. Averaging the instantaneous active force on the filament, we find that the average active force *per* motor *F*_*m*_ is related to the average stretch of the motors ⟨*y*⟩ by

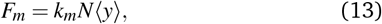

where *N* ∈ (0, 1) is the average fraction of motors attached. The value 0 implies that all motors are detached, while *N* = 1 implies that all motors are attached. In Equation (13), motor activity is treated as an internally distributed force density per motor (mean). That is, we have replaced the group of active motors (attached and detached) in the element *ds* centered on *s* with a representative (idealized) motor with an average extension ⟨*y*⟩.

In the biologically relevant over-damped limit where inertia may be ignored, force balance dictates that *u* satisfy the equation

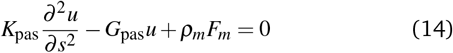

where *K*_pas_ = *Ew* and *G*_pas_ = *ρ*_N_*k*_N_. In the absence of activity *F*_*m*_ = 0, and Eqn. (14) predicts that local perturbations in *u* decay exponentially with a length scale 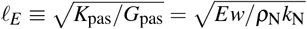 indicating, as expected, the crucial role played by the elastic resistance of the assembly to shear.

### 3.2 Motor kinetics

In the mean-field limit, a set of population balance equations relate the attached and detached probability densities *P*_*a*_(*s, y, t*) and *P*_*d*_ (*s, y, t*) to the attachment and detachment fluxes *J*_*a*_ and *J*_*d*_. These equations involve transition rates *v*_on_(*y*) and *v*_off_(*y*) between the two possible motor states and are

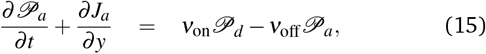

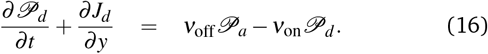

In our analysis, we treat the attachment frequency *v*_on_ in (15) and (16) as constant; while the probability of detachment is allowed to vary with motor extension *y*(*s, t*). Thus, we write *v*_off_ = *v*_off_(*y, δ*_*m*_), where *δ*_*m*_ is the characteristic motor extension at which the detachment rate becomes significant. Here, *d*_*m*_ and *δ*_*m*_ are each related to motor properties and kinetics and since we have ignored Brownian diffusion *J*_*a*_ = (*dy/dt*)*P*_*a*_.

### 3.3 Averaging and mean-field equations

Consistent with previous studies ^9,10^, we specify that the detached motors relax almost instantaneously to an equilibrium state with mean detached density *ρ*_*d*_ and zero motor extension. Simultaneously we assume ^9^ that the extension of attached motors is sharply peaked about ⟨*y*⟩, thus writing *P*_*a*_ = *ρ*_*a*_(*t*)*δ* (*y* − ⟨*y*⟩). where *δ* is the Dirac delta function. The Dirac delta function allows us to replace *v*_off_ and *v*_off_ in equations (15) and (16) with the attachment rate (mean-field) 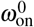 and the detachment rate (mean-field), ⟨*v*_off_⟩ = *ω*_off_. Using these relationships and integrating Equation (15) over *y*, we obtain the equation for the density of attached motors,

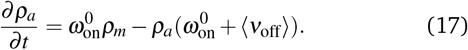

The mean field attachment frequency in (17) is assumed to be independent of motor strain. The form of the detachment rate *ω*_off_ is chosen to depend on the mean strain ⟨*y*⟩. In our model for active motors, the detachment rates increase with motor strain. Following previous work ^10^, we assume a characteristic motor extension *δ*_*m*_ beyond which detachment is preferred. The quantity 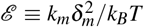 provides a strain-based measure of the energy needed for motor detachment. Based on this picture, we set 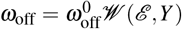 where *Y* ≡ ⟨ *y*⟩*/δ*_*m*_. Alternate forms of the function yield qualitatively similar results, provided that *W* is continuous and increases monotonically with *Y*.

To obtain a closed set of averaged equations, we need an expression for the evolution of. ⟨*y*⟩. We anticipate that this expression will incorporate three effects - the pre-extension of attaching motors, the fact that motors detach at finite extension, and the evolution of the extension whilst the motor is attached. Multiplying (15) by *y*, then integrating over *y*, and finally using ⟨*v*_off_ *y*; ⟩≈ ⟨*v*_off_ ⟩ ⟨ *y* ⟩ we find

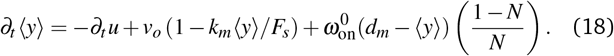

In Equation (18), the first term on the right hand side is the stretch due to the passive motion of the attached motor with *u* satisfying Equation (14)). The second term gives the motor velocity relative to the filament. The third term corresponds to the rate at which the mean strain changes due to the kinetics of motor attachment and arises from the difference in the extension values of attaching motors and detaching motors.

**Table 1.**
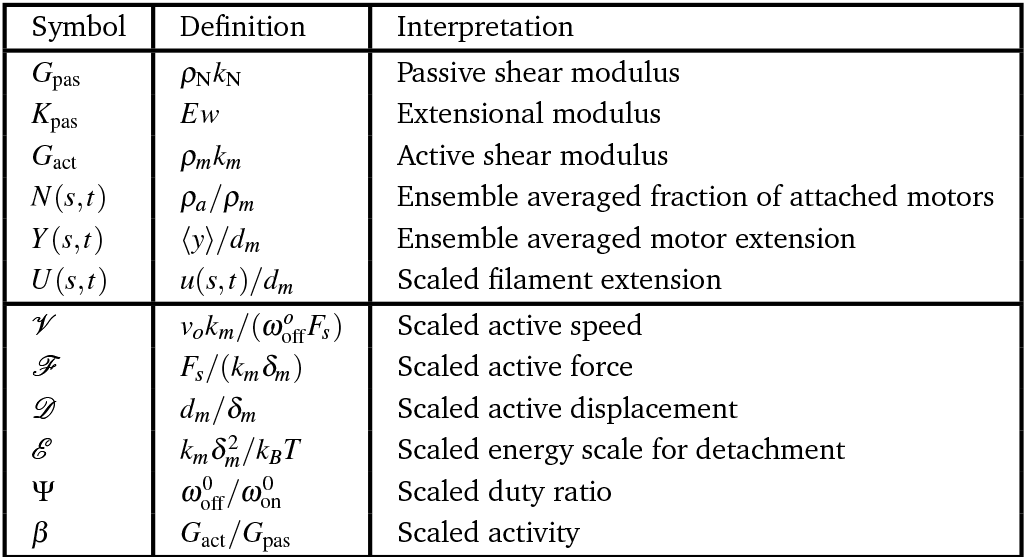
Elasticity parameters and dimensionless parameters that appear in the mean-field model – Eqns. (14), (17), (18), (19) and (21).

Equations (14), (17), and (18) constitute our model-field noiseless model. Scaling time with 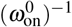, displacements by *δ*_*m*_, and defining scaled variables *ρ*_*a*_(*s, t*)= *ρ*_*m*_*N*(*s, t*) and *U* = *u/δ*_*m*_, we obtain dimensionless forms of Eqns. (17) and (18)

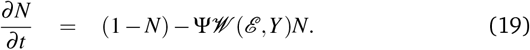

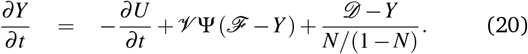

Equation (20) highlights that changes in the scaled motor displacement (strain) *Y* occur due to three effects: passive convection of the motor head when attached, the pre-extension of attaching motors, and additional effects that arise due to the fact that motors detach at finite extension.

Finally, we combine (12), (13) and (18), and rescale the length using 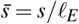 to obtain

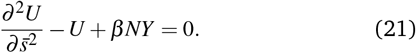

Here *β* ⟨ *G*_act_*/G*_pas_ where *G*_act_ = *ρ*_*m*_*k*_*m*_ is the active analogue of the passive shear modulus. Equations (19)–(21) constitute our mean field model and involve four dimensionless parameters: (1) the scaled active speed (quantifying motor speed at zero load) 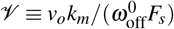, (2) the scaled active force ℱ ≡ *F*_*s*_*/*(*k*_*m*_*δ*_*m*_), (3) the scaled motor pre-strain *D* ≡ *d*_*m*_*/δ*_*m*_, and (4) the kinetic parameter 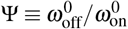 that is related to the motor duty ratio.

Our mean field approach leading to equations (19)–(21) complements and differs from previous attempts in important ways. First, consistent with experiments suggesting that bond failure is more naturally dependent on extension and only weakly on the extension rate, we have chosen 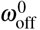 to depend on motor exten sion^9,10^ and not the extension rate ^55^. Second, non-linear coupling between passive and active deformations in (17)-(21) distinguishes our model from previous studies of motor-mediated bending of filaments ^55^. Finally, our model filament is weakly extensible, bolstered by recent experimental evidence ^3,7,18^. We have also ignored any viscous drag forces due to ambient fluid (see ESM §2 for a discussion of the role of viscous drag). We note that *V*, ℱ, *D* and Ψ appear in Eqn. (21) through a term multiplied by *β*.

### 3.4 Predictions of mean field model

Equations (19)–(21) permit a unique steady homogeneous solution. To find this we set *∂U/∂t* = *∂Y/∂t* = *∂ N/∂t* = 0 to obtain

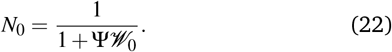

Here, *W*_0_ ≡ *W* (*Y*_0_), and 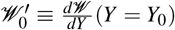. The steady-state extions is

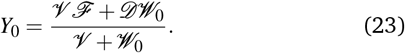

#### 3.4.1 Instabilities of active rigid segments

Before we study the strain field for an extensible filament, we first examine the stability of stationary solutions for a small rigid fragment with a length much shorter than the relaxation length of pure extensions, i.e. 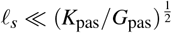. In this limit, we can ignore the extensibility of the segment. The dynamics of this segment relative to its neighbors can be mapped to that of a homogeneous population of motors acting on a rigid segment and working against an external spring with effective stiffness *K*_s_ ∝ *G*_pas_. The effective spring constant may be treated as an overall measure of the extensional resistance that also has active contributions arising from the attached motors in neighboring filaments. If *ρ*_N_ = 0, then the passive part of *K*_*s*_ ∝ *K*_pas_. Since the segment is inextensible, Eqn. (21) becomes

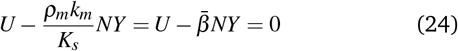

For *K*_*s*_ *>* 0, a translating filament is nonphysical and, hence, we probe the existence of oscillatory solutions. To do this, we analyze the stability of the base state given by Equations (19), (20), and (24) for small changes in the fields given by *N* = *N*_0_ + *N*_1_ exp(*σt*) and *Y* = *Y*_0_ + *Y*_1_ exp(*σt*) where *N*_1_ ≪ *N*_0_ and *Y*_1_ ≪ *Y*_0_. Linearizing equations about the base state (*N*_0_,*Y*_0_), we recast the resulting equations in matrix form and analyze the structure of the eigen-values in two limiting cases.

**Case 1:** When *V* = 0 and *D* ≠ 0, the neutral stability curve is given by

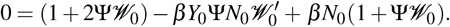

The frequency of emergent oscillatory solutions is

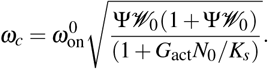

In Figs. 3(a)-(c) we show solutions to equations (19), (20) and (24) for a subset of parameter values. These illustrate the steady expression for the critical frequency at onset, and stable oscillatory dynamics exhibited by the motor-driven rigid segment. In Figure 3(a-i), we plot time-dependent solutions obtained using a standard Runge-Kutta (RK4) method for parameters that result in stable oscillations. Since the parameters correspond to an operating point slightly displaced from the neutral stability curve, the resulting stable solution is impacted by nonlinear terms in Eqns. (19) and (20) and resembles relaxation rather than sinusoidal oscillations. The blue curves correspond to Ψ = 0.3 while the red curves correspond to Ψ = 1.0. The activity parameter, that is, the scaled motor density 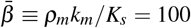, and we have set *D* = 1.0, *Y*_0_ = *D, V* = 0, and *W* = exp(2|*Y*|). For comparison (see also ESM §1), in Fig. 3(a-ii), we illustrate the nature of solutions when viscous drag due to an external Newtonian fluid is considered. As derived in ESM §1, Eqn. (24) is then modified by adding a term proportional to the drag parameter *A* which is here set to 2.0. Other parameters are *D* = 1, and *V* = 0. We fix 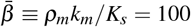 and Ψ = 0.5. The blue curves are for *W* = exp(2|*Y*|), and the red curves correspond to *W* = cosh(2*Y*).

**Fig. 3.**
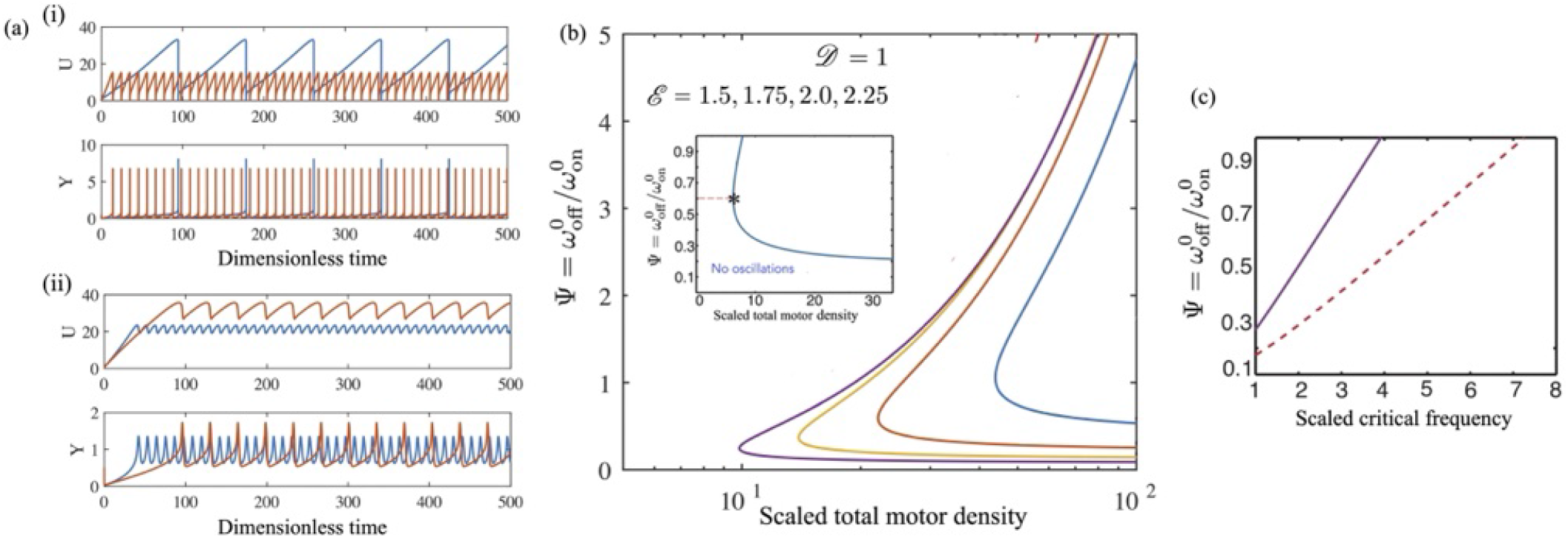
Steady states, and oscillatory dynamics of a small rigid segment animated by molecular motors working against a resisting spring modeled by equations (20) and (24). (a) Typical oscillations in time *T*, with the periodic functions (here non-linear, and in the linearly unstable region away from the critical point) demonstrating features seen in relaxation oscillations. (i) Results when the drag due to the external fluid may be neglected. The blue curves correspond to Ψ = 0.3 while the red curves correspond to Ψ = 1.0. Here, 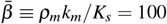, *D* = 1.0, *Y*_0_ = *D, V* = 0, and *W* = exp(2|*Y*|). (ii) Oscillations when the external fluid is Newtonian. Here, *A* = 2.0, *D* = 1, *V* = 0, 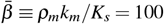 and Ψ = 0.5. The blue curves are for *W* = exp(2|*Y*|), the red curves correspond to *W* = cosh(2*Y*). (b) Neutral stability curves that separate the steady stable solutions from stable oscillations for negligible external fluid drag. Here, parameters are *D* = 1, and *V* = 0. The curves here correspond to *W* = exp(ℰ |*Y* |) – these may be compared to ESM Figure-3 where we compare the results here to the case when *W* = cosh(ℰ*Y*). From left to right, parameter ℰ = 1.5, 1.75, 2.0 and 2.25. (Inset figure) Here we focus on the neutral stability 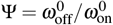 vs. the scaled motor density *ρ*_*m*_*k*_*m*_*/K*_*s*_ when 0 *<* Ψ *<* 1 and for large ℰ (here = 4.944). The pre-strain value *Y*_0_ = 0.5. We observe a gentler curve around the turning point. (c) Scaled frequency of oscillations at the critical point, 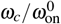, indicated with the ⋆ symbol in inset figure 3(b), as a function of Ψ. The solid line is the exact result while the dashed line is the adiabatic approximation.

In Fig. 3(b), we show neutral stability curves that separate steady stable solutions from stable oscillations in the absence of external fluid drag. Here, the chosen parameters are *D* = 1, and *V* = 0. The detachment function investigated here has the form *W* = exp(ℰ |*Y* |) with ℰ = 1.5, 1.75, 2.0 and 2.25. This figure may be compared with ESM Figure 3 showing the neutral stability curve for *W* = cosh(ℰ*Y*). We note that for each value of ℰ, there is a critical value of Ψ(ℰ) below which the base state is linearly stable and there are no oscillations. In contrast, for a fixed value of *β*, the oscillations are stable for a range Ψ_L_(*β*) *<* Ψ *<* Ψ_U_(*β*). In the inset, we focus on the region close to the turning point and examine the dependence of the neutral stability curve 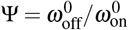 vs. scaled motor density, *ρ*_*m*_*k*_*m*_*/K*_*s*_ when ℰ = 4.944. To illustrate how both the form of *W* and the value of ℰ determine the curvature near the turning point, we show the neutral stability curve over the range 0 *<* Ψ *<* 1 when *W* = cosh(ℰ*Y*). For fixed Ψ, a critical number of motors is required for the oscillations to manifest (the value of Ψ = 0.6 is indicated as an example). In general, the critical motor density depends on Ψ, and on *K*_*s*_*/*(*ρ*_*m*_*k*_*m*_).

Finally, in Fig. 3(c), we plot 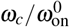, the (scaled) frequency of oscillations, as a function of Ψ. The solid line is the exact analytical result presented in the ESM. The dashed line is the adiabatic approximation where motor extension is slaved to filament dynamics. A representative critical point is indicated with ⋆ (inset of Fig. 3(b)). Analytical expressions for the neutral stability curves (ESM §I), predict that the neutral stability curve depends on Ψ, ℰ and *β*.

Incorporation of viscous drag due to the viscosity of the external medium modifies the neutral stability curve and the value of the emergent frequencies, but otherwise does not change the results qualitatively (ESM §1). Detailed exploration of the boundary that separates the static homogeneous solution from stable oscillations shows that stable oscillatory solutions exist only within a certain range of Ψ.

**Case 2:** Next, we consider the complementary limit ^10^ *D* = 0 and *V* ≠ 0. In this case, linear stability analysis delivers the expression for the critical frequency at onset,

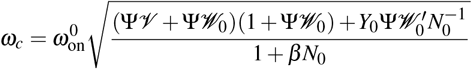

and the equation for the neutral stability curve,

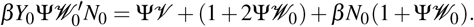

When motors are adiabatically coupled to the dynamics of the segment so that motor kinetics is always faster than the sliding velocity, 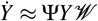, and the critical frequency reduces to

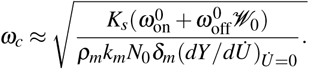

Overall, our results suggest that oscillations are driven by positive feedback, that is, the emergence of negative effective spring constants and/or negative friction coefficients that arise from the interplay between motor kinetics and external elasticity. For large values of Ψ, the number of motors attached is insufficient to supply the energy for the oscillations, suggesting an upper bound. For small Ψ, too many attached motors increase elastic resistance and friction, causing a strong damping of oscillations. In the stable oscillating state, the input of active energy to the system balances the irreversible viscous dissipation caused by motor friction.

#### 3.4.2 Persistence length for assemblies in rigor

An important case to consider is when active assemblies are in rigor, that is, there is no intrinsic motor activity. This could happen due to chemical quenching of motor activity such as, for example, decreasing ATP or Ca^2+^ levels to values that hinder motor activity. The motors are then stuck in fixed configurations (attached or detached), and therefore we have *ω*_on_ = 0, *ω*_off_ = 0 and *v*_*o*_ = 0. In this limit, the motors are permanently bound to the filament track and can actually sense the instantaneous displacement of the attachment point (where the head is) rather than just the rate of displacement. The effective elasticity of the active assembly may then be estimated as follows. The number of attached motors *N* cannot change as the attached motors cannot be detached. Let *N* = *N*_0_ be the pre-existing attached fraction. The motor extension *Y* responds passively to the change in the attachment point of the motor head, so that

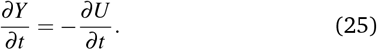

Integrating this equation and setting *U* (*t*) −*U*_0_ = *δU*, we get

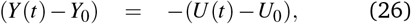

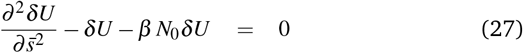

from which we deduce that the characteristic persistence or decay length scale λ_R_ in the rigor state is

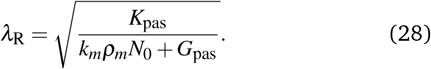

The effective shear modulus due to the links, *G*_eff_, is obtained by equating the force needed to shear the composite - here, the motor springs act as passive springs. This procedure yields (*N*_*R*_*ρ*_*m*_*k*_*m*_ + *ρ*_*N*_*k*_*N*_)(*ds b*)⟨*y*⟩ = *F*_*s*_ = *G*_eff_(⟨*y*⟩*/D*)*ds b*, thus yielding an estimate for the effective modulus

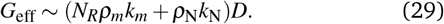

#### 3.4.3 Persistence lengths for assemblies out of rigor Steady strains

From (21), we note that stationary, steady filament displacement, *U*_0_, attached motor fraction, *N*_0_, and motor strain, *Y*_0_ fields satisfy

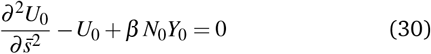

where ℱ_0_ = ℱ (*Y*_0_) with *N*_0_ and *Y*_0_ given by equations (22) and (23). We choose boundary conditions consistent with the end corresponding to 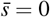 fixed so that *U*_0_(0)= 0 and with the end corresponding to 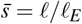 left free and satisfying 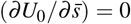. Solving Equation (30) with these boundary conditions, we find

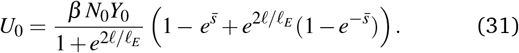

The strain *U*_0_ is therefore linear in *β*, and surprisingly, the decay length is independent of *β*. This result may be rationalized as follows. Attached motors sense the strain rates of the filament but do not have any way to gauge the amplitude (magnitude) strain at the point of attachment at the time of attachment or later. Passive crosslinkers, on the other hand, sense both absolute strain and strain rates. Extending the analysis to the case where *ρ*_*m*_ is spatially varying (inhomogeneous total motor density), we find that although persistence lengths can vary locally and deviate from *ℓ*_*E*_, it is still independent of *β*.

##### Oscillatory strains

The final question we address in the context of the continuum model is the decay of oscillatory strains in a long filament. Oscillations in biology typically have sharply defined frequencies; thus resulting deformation fields may also have well-defined frequencies, an example being ciliary oscillations. Here, one can imagine there being two or more oscillating local regions in a long flagella and ask if and when these regions can mechanically sense each other via strain or deformation fields. Such interactions are possible if these regions are within a decay length of each other.

We now investigate how localized oscillatory displacements (imposed at one end or naturally emergent within the filament) and associated extensional strains decay with distance. To derive estimates of the decay length, we consider an undeformed filament that has oscillations imposed at *s* = *ℓ*, while still respecting the constraints that lead to (31). This is achieved, in an asymptotically accurate fashion, by subjecting the free end to a small-amplitude oscillatory displacement. This oscillation may correspond to an externally imposed perturbation or an intrinsic emergent oscillation within a filament, as analyzed in § 3.4.2. To allow a homeostatic response to emerge, the frequency of the oscillation must be sufficiently small. Since oscillatory instabilities close to onset have Fourier forms, we consider boundary conditions

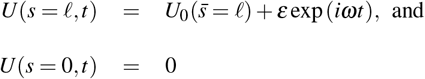

with *s* ≪ 1. The homeostatic response implies that the strain and motor fields (motor density and mean motor stretch fields) along the whole filament eventually oscillate at the same imposed frequency *ω*. However, the fields *U* (*s, t*), *N*(*s, t*) and *Y* (*s, t*) will have different amplitudes and spatially variable phase-lag components.

Of interest is the long-time response, as *t* → ∞ after the initial transients have decayed. The solutions then must have the form

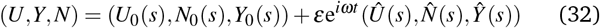

where the terms 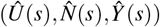 incorporate phase-lag variations. Note that we require *Ŷ*(*s* = *ℓ*) to be perfectly in-phase with the imposed forcing. Substituting (32) into equations (19)–(21), and considering only *O*(*ε*) linear terms, we find that the disturbance strain field *Û* (*s*) satisfies

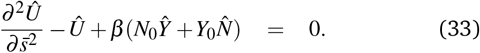

Linearized equations for motor dynamics have the form

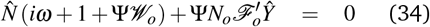

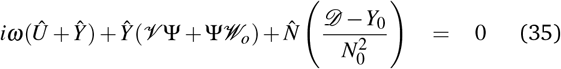

from which we can derive an effective viscoelastic response function 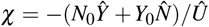. The function *χ* depends on the stall force *F*_*s*_, the free velocity (motor speed at zero load *v*_0_) and the duty ratio Ψ (see ESM §2.2). Solutions 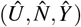, for oscillations imposed at *s* = *ℓ* with the filament fixed at *s* = 0, are obtained by solving equations (33)–(35) with boundary conditions 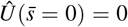 and 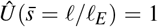. With *χ* determined for the oscillatory forcing at the end *s* = *ℓ*, Equation (33) can be recast as

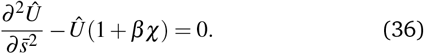

Let us first consider the special case of very long filaments. It is convenient to redefine the variable *x* = (*ℓ* − *s*) so that we can invoke the limit *ℓ/ℓ*_*E*_ → ∞ and set the boundary condition as *x* → ∞. we seek decaying solutions to Eqn. (36) of the form *Û* = *Û*_∗_ exp(−*qx*). For such solutions to exist, consistency requires that

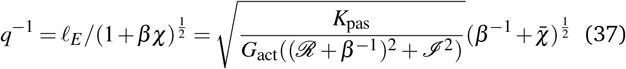

where 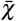 is the complex conjugate of *χ* and we have decomposed *χ*(*ω*) ⟨ ℛ (*ω*)+iℐ (*ω*) into its real and imaginary parts. ESM §2.2 extends this analysis to the full range of *ℓ/ℓ*_*E*_. In general, the values of *q* required from consistency and solvability are imaginary. The length scales over which the amplitude of the strain field decays and the scale over which the field varies are

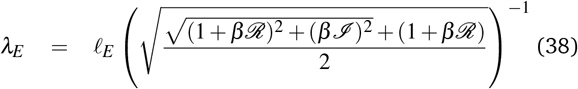

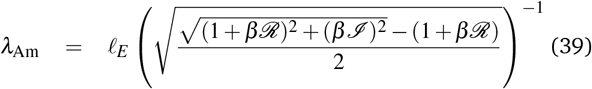

motivating the general form

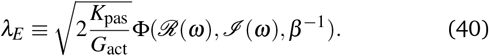

Eqn. (40) indicates that the persistence length is a product of two terms that encapsulate different physics. The frequency independent term 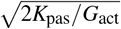 combines the passive extensional modulus of the filament with the active shear modulus due to the attached motors. The function Φ incorporates multiple effects. First, it incorporates shear stiffening due to permanent passive linkers through *β*. Furthermore, it also incorporates frequency dependence and motor kinetics through the term *χ*. Analytical expressions for the function *χ* are presented in the ESM, with explicit relationships highlighting the dependence of Φ on *A*_1_, *A*_2_, and *A*_3_ outlined (ESM, equations (35), (36) and (43)). Furthermore, focusing on the limiting case where *A*_1_ = 0, we obtain analytical expressions for the real and imaginary parts of *χ* in two limits (ESM, equations (38)–(41)): *ω* ≫ *τ*^−1^ and *ω* ≪ *τ*^−1^ where the duty ratio *τ* ⟨= *τ*_*a*_ */*(*τ*_*a*_ + *τ*_*d*_)= 1*/*(1 + Ψ*ℱ*_0_). Here, *τ*_*a*_ and *τ*_*d*_ are the load-dependent attached and detached times averaged over a motor mechanochemical cycle. From Eqn. (38) we also deduce the existence of a crossover value of *G*_pas_ that separates the active motor dominated regime from the passive linker dominated regime. The pre-strain *U*_0_ sets the base state properties that determine the frequency dependence of *χ*. This has implications for the elasticity-mediated communication between localized oscillating regions on the filament.

Although our mean-field theory provides a baseline understanding of the underlying physics, the analysis has limitations. First, it does not shed light on what happens when the number of motors is small when a continuum model may not be the best approach. Second, the mean-field model ignores athermal noise arising from finite motor number addressed next.

## 4 Brownian-MPC simulations of noisy assemblies

Although thermal (Brownian) noise can be neglected in the case of stiff filaments, a second type of noise, *discrete noise*, can arise due to the binding and unbinding of a finite number of motors to the filaments. This athermal stochastic noise can have significant effects on the filament dynamics, especially when at low motor densities ^59–61^. In the context of flagella, the onset of steady or oscillatory extension necessarily involves a small number of motors when the deformation field develops from intially noisy states.

### 4.1 Discrete model

Having studied the role of shear stiffness due to passive links in controlling the bending and extension of deformed filaments, we now describe our discrete, microscopic model that includes noise due to motor activity. The main aim of the discrete simulation is to qualitatively compare the influence of noise in systems containing filaments and motors by selecting a finite number of motors. Additionally, since we can also adjust the discrete version of filament elasticity, we can also study moderate or soft filaments, regimes that are inaccessible to mean-field theory.

The discrete analog of the weakly extensible filament is modeled as a one-dimensional chain of *M* rigid segments. Each of these segments is made up of *M*_*m*_ discrete spherical monomers of diameter ∀ located at *r*_*i*_,(*i* = 1,…,*N*_*m*_), connected by an elastic potential

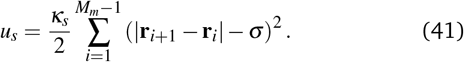

Each segment has length *ℓ*_*s*_ = 80 *σ* and is connected to neighboring segments by linear elastic springs of stiffness *K* with rest length 80*σ*. The first segment is connected to a rigid wall by the same elastic potential at the location corresponding to *s* = 0 while the other end of the compound at *s* = *ℓ* is kept free. Bending stiffness is implemented via a three-body bending potential

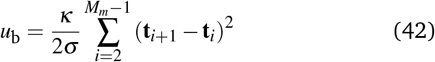

where **t**_*i*_ = (**r**_*i*_ − **r**_*i*−1_)*/*|**r**_*i*_ − **r**_*i*−1_ |. Here, effective bending rigidity *κ* penalizes angular changes from a local straight geometry for the segments. We keep the bending elastic constants *κ*_*s*_ = 2 ×10^4^(*k*_*B*_*T/σ*^2^) and *κ/σ* = 2 × 10^4^ (*k*_*B*_*T*) with an appropriately large persistence length for the segment ~ 250 *ℓ*_*s*_. This ensures that the segments are practically rigid and inextensible, although the extensibility of the entire filament can still be tuned by adjusting the elasticity parameter *K*. Although our model can be used to study both elongation and bending deformations by treating *κ* as a variable parameter, here we focus solely on the extensional deformation of the filament.

Motors imposing active forces that cause filament shear and extension are modeled as linear elastic springs of stiffness *k*_*m*_ and rest length *ℓ*_*m*_. These motors are uniformly distributed on each segment with a number density *ρ*_*m*_. One end of the motor (tail) is grafted to a point **r**_*t*_ on a rigid base, while the other end (head) can attach and detach to a given monomer, located at **r**_*h*_, of the filament. During the attachment process, the monomer bead that lies closest to the motor head is chosen so that the motor extension, |**r**_*h*_ − **r**_*t*_ | − *ℓ*_*m*_, upon attachment is minimal. Similarly to the mean-field model, we assume a constant attachment probability *p*_on_ for each motor. Once attached, the motor head steps to the next monomer in the direction of the anchored end, *s* = 0, with a discrete step size equal to the diameter of the monomer *a*. As motor heads move, the accompanying extension of motor length leads to a force **F**_*m*_ = −*κ*_*m*_(|**r**_*h*_ − **r**_*t*_ | − *ℓ*_*m*_) **ê**, where the unit vector **ê** = (**r**_*h*_ − **r**_*t*_)*/*|**r**_*h*_ − **r**_*t*_ |, which acts on the attached filament. This force is balanced by the tension on the filament to which the motor is attached, thus resulting in localized active strains.

Finally, we assumed a load-dependent stepping velocity of the motor head, choosing a linear relationship, *v*_*h*_ = *v*_*o*_(1 − |**F**_*m*_|*/ f*_max_) with *f*_max_ being the stall force *F*_*s*_ consistent with the form assumed in the mean-field theory and experimental observations. Motor heads that are attached to the filament sense the load and detach at a rate given by the load dependent probability 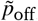, defined by the piecewise function

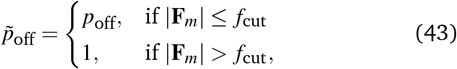

where *p*_off_ is the load independent detachment rate of motors and *f*_cut_ is the motor cut-off load. We take *f*_cut_ = *f*_max_ in our simulations, although in general they are not necessarily equal.

The load dependence of the detachment probability of individual motors allows for crucial two-way coupling between filament elasticity and the motor response and also sets the critical motor extension at which detachment occurs. Furthermore, the constant load-independent detachment probability *p*_off_ together with *p*_on_ produces an equilibrium fraction of attached motors, *p*_on_*/*(*p*_o*n*_ + *p*_off_). We recall that in continuum theory, the steady-state fraction of the attached motors is 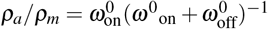. In the absence of any mean motor deformation (that is, when *Y*_0_ = 0 so that *ℱ* = 1), this fraction equals 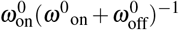 providing the mapping between the continuum theory and the stochastic simulations.

Despite being a discretized model, the segments in the filament still represent a coarse-grained description of the filament in biological systems, and hence we allow multiple motor heads to occupy the same site on the filament. Because of the initial distribution of the motors, the torque on the filament due to motor activity is negligible, and the system is essentially one dimensional.

For simplicity and to focus on the role of activity and noise, we did not incorporate permanent passive crosslinkers of the mean field model. Although our simulation corresponds to the limiting case *ρ*_*N*_ = 0, our results are general. This is because the shear resistance is always provided by the active motors when attached. With this simplification, the passive elasticity of the filament is solely contributed by the weak extensional elasticity of the filament.

The time-dependent evolution of this motor and segment system is analyzed in the overdamped limit using the Brownian multiparticle collision dynamics method (Brownian-MPC) ^62^. This method neglects long-range hydrodynamic interactions between the segments, while keeping the system in a thermal bath. According to this scheme, each monomer bead of the segment independently performs a stochastic collision with phantom fluid particles with momenta taken from the Maxwell-Boltzmann distribution with variance *ρk T*, where *ρ* is the density of the phantom fluid. In between collision events, the monomer position and velocity are updated using standard molecular dynamics (MD).

The time scale for Brownian MPC simulation is 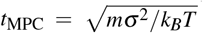, where *m* is the mass of a phantom-fluid particle and *a* is the diameter of the monomer bead that is equal to the length scale ^62^. However, the physical time scale relevant to this problem is the numerically calculated thermal relaxation time of the segment of length 80 *a*, which is much larger than *t*_MPC_. This re-laxation time is determined by the viscosity of the medium set by the simulation parameters ^62,63^. For the same set of fluid parameters used in reference ^63^, we find that the relaxation time, *τ*, of the length segment 80*σ* is approximately 4.6 × 10^3^*t*_MPC_.

### 4.2 Oscillations of a rigid segment

As in our mean field theory, we first study the dynamics of a single rigid segment of the long filament driven by an aggregate of active motors working against a resisting anchor spring to compare with the theoretically analyzed dynamics for filament fragments of length 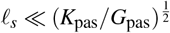. This allows us to characterize the dynamical regimes of that segment as a function of motor activity and elastic stiffness. More importantly, this allows us to identify the appropriate base state to use in our calculation of the decay lengths of imposed or inherent local extensional strains.

The discrete segment of length *ℓ*_*s*_ = 80 *σ* is forced by 800 motors uniformly distributed along its length (corresponding to the schematic in Fig. 4 (a). The end corresponding to *s* = 0 is anchored to a point using a linearly elastic spring (anchoring spring in Fig. 4(a)) of stiffness *K*_*s*_. The spring has, without loss of generality, a zero rest length. The dynamical response of this system is studied as a function of two parameters, a stiffness parameter *K*_*s*_*/k*_*m*_ and the activity parameter *p*_off_*/p*_on_. This is done by tracking the displacement of the segment as a function of time as well as the temporal variations in the fraction of active motors attached to the segment. Times are scaled using *τ* ≃ 4.6 × 10^3^*t*_MPC_. Depending on the values of *K*_*s*_*/k*_*m*_ and *p*_off_*/p*_on_ we find three distinct dynamical regimes, as illustrated in Fig.4(b). For a fixed value of *K*_*s*_*/k*_*m*_ when *p*_off_*/p*_on_ ≪ 1 most of the motors are attached to the filament, subjecting the segment to a significant active force. Consequently, the segment displays a steady extension.

**Fig. 4.**
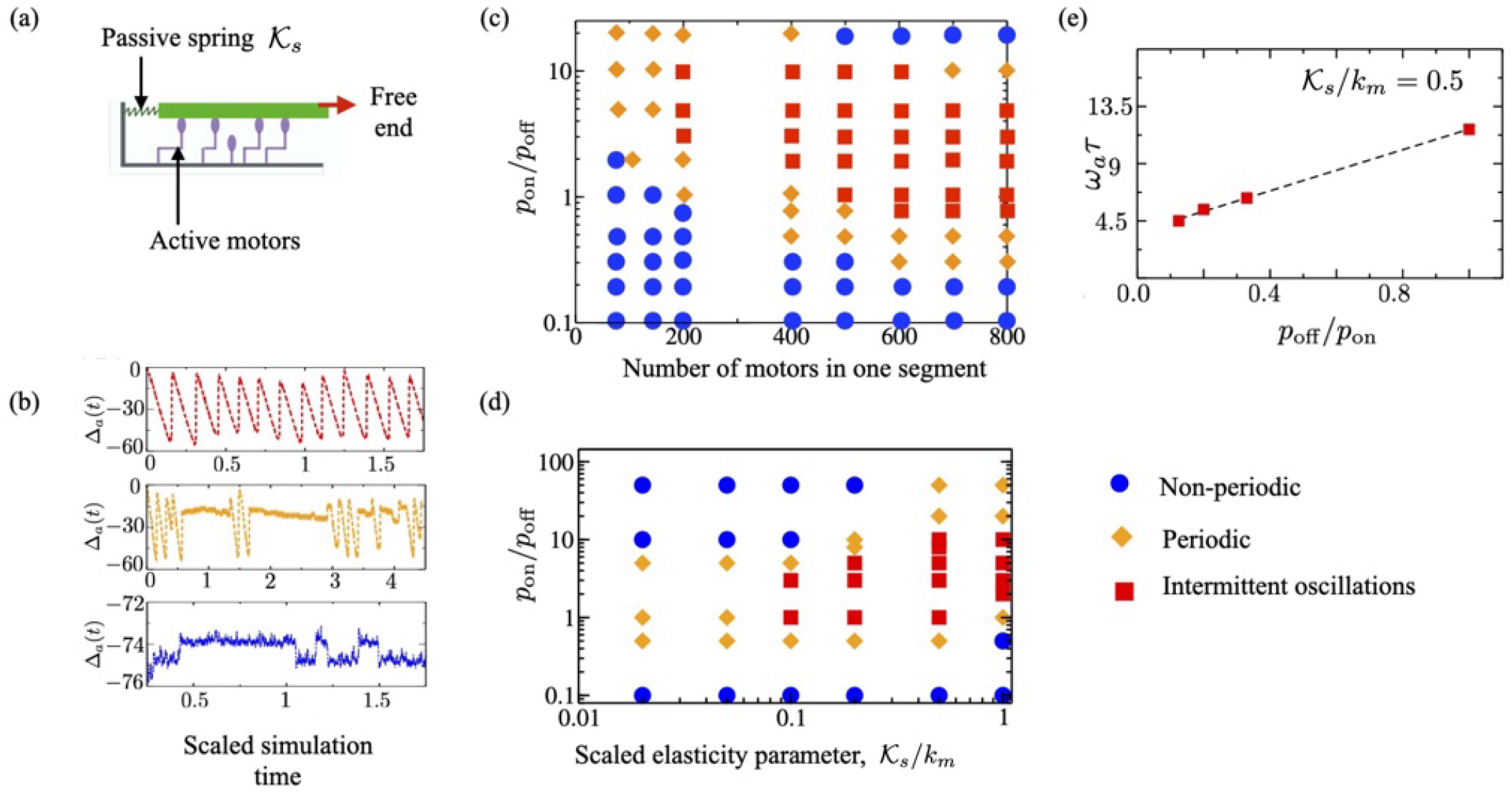
(a) Sketch of the filament-motor system for a rigid segment. Note that in the simulations we do not include any permanent passive linkers that attach the segment to the base. Rather, the rigid segment moves due to active forces imposed by attached motors (green) and this (translational) motion is resisted by passive forces here generated by an effective (anchoring) spring with stiffness *K*_*s*_. (b) The time dependent displacement, Δ_*a*_(*t*)*/σ* of the filament for a fixed ratio *K*_*s*_*/k*_*m*_ = 0.2, and for zero-load motor detachment/attachment probabilities, *p*_off_*/p*_on_ = 1.0 (top), 2 (middle) and 10 (bottom), illustrating various dynamical regimes observed in simulations. (c) Phase diagram from the Brownian-MPC simulations indicating various dynamical regimes mapped in terms of *K*_*s*_*/k*_*m*_, and *p*_off_*/p*_on_. Dynamical regimes are, steady extension (′), intermittent oscillations (lozenge), and steady oscillations (square). (d) The oscillation frequency of a periodically oscillating filament with *K*_*s*_*/k*_*m*_ = 0.5 as a function of *p*_off_*/p*_on_.

In the complementary limit, where *p*_off_*/p*_on_ ≫ 1 the reattachment rate of the motors is small, causing most of the motors to be in the detached state. Thus, in the absence of any significant extensional force, the filament attains a steady, but non-extended state. However, a third time-dependent regime is observed for intermediate values of *p*_off_*/p*_on_; in this regime, we observe intermittent (Fig. 4(b)-middle) or regular (Fig. 4(b)-top) oscillations in the segment extension. These oscillations are quantified by measuring the distance of the free end of the filament from the anchor point (Δ_*a*_) as a function of time. Stable oscillatory states have a well-defined frequency. In the intermittent oscillatory state, the segments switch randomly between stationary and oscillatory states.

The transition to an oscillatory state from a stationary state is consistent with predictions of our mean-field theory. A sweep of the parameter space yields the computational phase plot that we find to be qualitatively consistent with the theoretical one, cf. Figs. 3(b) and 4(c). For a fixed number of *total* motors equivalent to a fixed stiffness parameter, oscillations are observed only for a range of Ψ, which is, in turn, equivalent to a range of *p*_on_*/p*_off_ (or equivalently the inverse). In the absence of noise, the boundary in the parameter space between stationary and oscillatory states is sharp. Discrete noise, however, smoothens this sharp boundary and renders it fuzzy, thus the intermittent states. In the absence of noise, the boundary in the parameter space between stationary and oscillatory states is sharp. Thus, the intermittent states are sustained by noise - here due to the discrete motor binding and unbinding events and due to the finite number of motors. We emphasize that there is no thermal noise in the system and anticipate that the presence of thermal noise will probably broaden the intermittent region further.

Focusing on the regular oscillatory regime, we also compute the frequency as a function of the *p*_off_*/p*_on_ at a fixed stiffness ratio as shown in Fig. 4(e). We find that the response is consistent with the mean-field predictions, with the frequency of oscillations *ω*_*a*_ increasing with the ratio *p*_off_*/p*_on_. This can be compared to Figure 3(c)-(ii) where the oscillation frequency is seen to increase almost linearly with increasing values of Ψ.

The physical picture underlying the onset of these oscillations may be understood as follows. The motion of the filament due to motor activity results in a non-linear viscoelastic coupling between the motors, the spring and the irreversible motor friction. At criticality, the motion is determined by the linear response to small perturbations. In this limit, the viscoelastic coupling may be reduced to effective, linear, in-phase (effective elastic) and out-of-phase (viscous) compliance terms. Oscillations arise when either of these terms turns negative as a result of positive feedback. When the system is in the stable oscillatory state, the power input to the system due to motor activity balances the energy dissipated due to frictional damping caused by motor attachment. Invoking expressions from the theory presented earlier, we estimate the active friction due to the motors to be 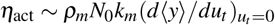 where *u*_*t*_ = *du/dt* and *ρ*_*m*_*N*_0_ is the total number of attached motors. As Ψ increases, *N*_0_ and 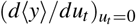 decrease, and the frequency of the oscillations increases.

### 4.3 Decay of steady strains

Having identified the dynamical regimes of a single rigid segment, we now analyze the dynamical behavior of a long, extensible composite filament (sketched in Fig. 5(a)). This composite filament is made up of 50 rigid segments, each of length 80*σ*. The softness of the filament is controlled by the elastic constant of connecting linear springs, *K*_*s*_ (∝ *K*_pas_ in the continuum theory). We distribute 80 motors uniformly on each segment and restrict parameters to lie in the regime where the filament undergoes a steady, non-oscillatory extension, *u*_0_(*s*). The phase plot shown in Figure 3(c) allows us to choose such appropriate parameters. Due to the stochastic nature of discrete motor binding and unbinding events, all steady quantities have fluctuating components. We measure the mean of these quantities by taking a time average after the system has evolved into a steady state.

**Fig. 5.**
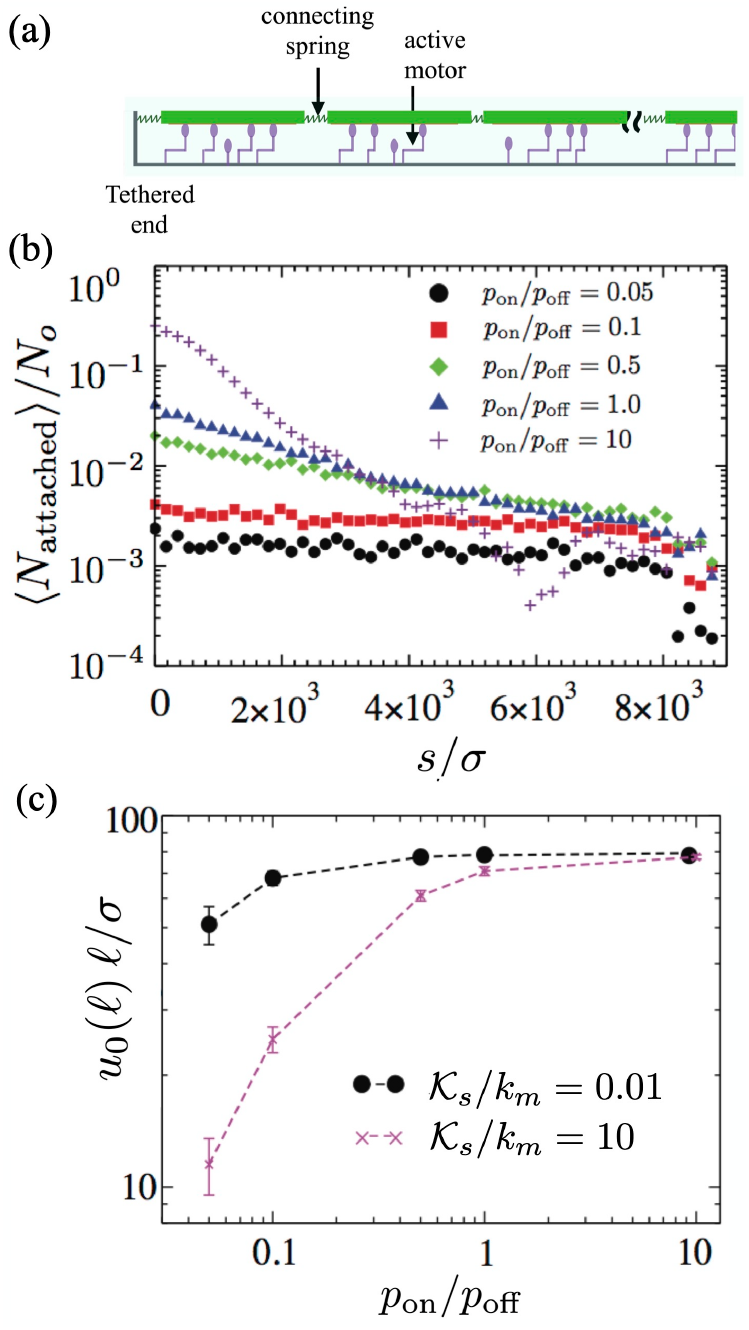
(a) Schematic of the discrete composite filament used in our sim-ulations. (b) The fraction of motors attached on the composite filament ⟨*N*_*a*_⟩, as a function of their position from the clamped end for *K*_*s*_*/k*_*m*_ = 10. The mean motor attachment is maximum close to the clamped end. (c) Time-averaged steady state extension of the segment at the free end, ⟨*u*_0_(*ℓ*) ⟨, as a function of *p*_on_*/p*_off_, for various values of passive spring stiffness, *K*_*s*_*/k*_*m*_. For large *p*_on_*/p*_off_ the extension saturates at a value set by the finite length of the composite.

Since motor kinetics is coupled to inhomogeneous filament strain, motor-imposed filament extensions modify the number of motors attached to the filament. In Fig. 5(b) we plot the mean fraction of motors attached, ⟨*N*_*a*_⟩, as a function of the distance from the anchored end, *s* for *K*_*s*_*/k*_*m*_ = 10 and for different values of the ratio *p*_on_*/p*_off_. When *p*_on_*/p*_off_ *<* 0.1 the motor-imposed extension of the filament is weak as the number of attached motors is small and the motor distribution is roughly uniform along the filament. However, when *p*_on_*/p*_off_ *>* 1, enhanced motor attachment imposes significant extension to the filament, which is maximum at the free end.

The non-uniform extension of the composite leads to a nonuniform distribution of attached motors with the mean motor attachment being maximum near the anchored end (*s* = 0), where the filament extension is minimum as seen in Fig. 5(c). The attached fraction decreases with the distance from that point as the filament strain increases as one moves towards the free end. This trend has implications when the total motor fraction is inhomogeneous. Lower total motor numbers near clamped ends, combined with expected lower attached motors as evidenced in Fig. 5(c) would then yield impactful changes to local strain gradients.

The time-averaged steady motor-induced extension ⟨*u*_0_⟩ of the filament, as a function of the stiffness parameter *K*_*s*_*/k*_*m*_ is shown in Fig. 6 for two values of the activity parameter *p*_on_*/p*_off_. Note that *p*_off_ is constant in these plots. Consistent with Fig. 5(c), filament extension is minimum near the clamped end and maximum at the free end; furthermore, both *p*_on_*/p*_off_ and *K*_*s*_*/k*_*m*_ equally influence the value and the gradient in the extensional strain. In general, *u*_0_ increases with the fraction of attached motors (∝ (*p*_off_*/p*_on_)^−1^) and the softness of the filament (∝ (*K*_*s*_*/k*_*m*_)^−1^). In the continuum limit, as anticipated by the mean field theory, the steady-state extension (*u*_0_) in the limit of no permanent cross-linkers is *u*_0_(*s*) ∝ (*ρ*_*m*_*k*_*m*_*/*2*K*_*s*_) *s*(2*ℓ* − *s*) where the constant of proportionality depends purely on motor kinetics and is independent of position (see ESM §3). In the absence of passive cross-linkers, *u*_0_(*s*) does not have any characteristic decay length; this is because the active motors form only temporary attachments and do not contribute to shear stiffening.

**Fig. 6.**
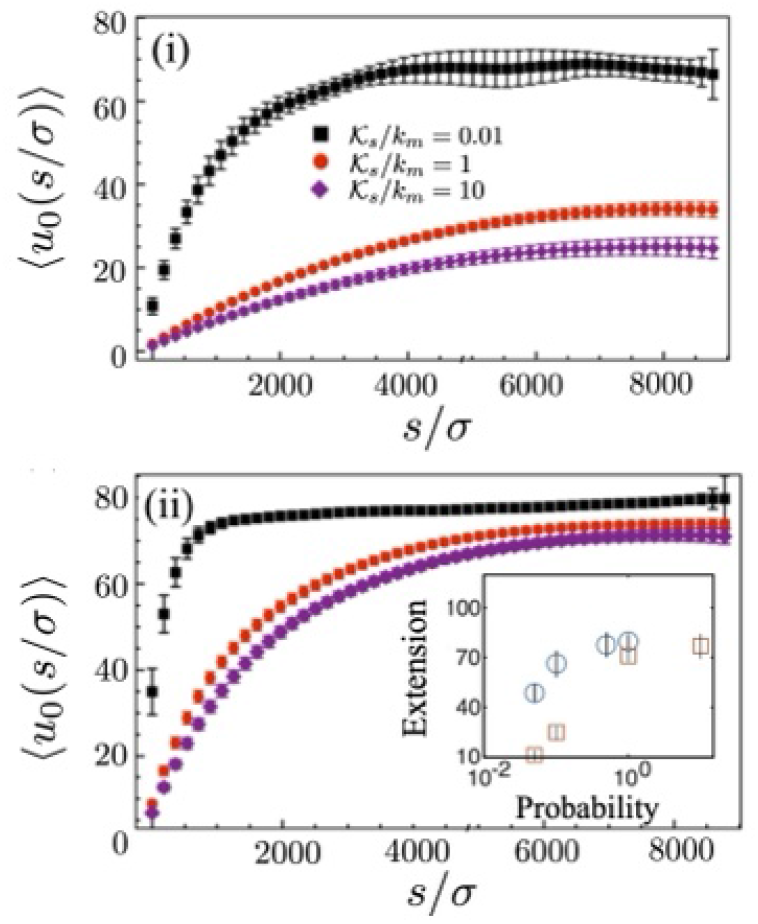
The dimensionless mean steady state extension of material points as a function of their position from the clamped end *s* = 0 for *p*_on_*/p*_off_ of (i) 0.1, and (ii) 1. Vertical bars correspond to the root mean square deviation. The extension is less correlated for soft assemblies (*K*_*s*_*/k*_*m*_ = 0.01) while it is highly correlated for stiff ones (*K*_*s*_*/k*_*m*_ = 10). The strain gradient localizes close to the clamped end for soft assemblies, especially for large attachment probabilities. We expect large fluctuations to have more of an effect in disrupting coherent behavior in these domains. Other parameters correspond to Figure 5. (Inset) Extension at the free end as a function of *p*_on_ for soft (*K*_*s*_*/k*_*m*_ = 0.01, circles) and stiff (*K*_*s*_*/k*_*m*_ = 10, squares) filaments.

The Brownian-MPC simulations are an improvement over the mean-field theory in an important way. In the theory, the local attachment fraction was spatially invariant and independent of position along the filament. In our case, the coupling between elasticity and motor kinetics is intrinsic and nonlinear. A consequence of this is the strong variation of the attached motor fraction with the position of the clamped end observed for high attachment probabilities. The quadratic nature of the extension field predicted from mean-field theory is qualitatively consistent with simulation results. By allowing for the attached motor fraction to vary with position and associated nonlinear coupling effects, we can explain the deviations from the quadratic scaling of the extensional strains closed to the clamped end seen in Figures 5(a)-(c).

The discrete nature of motor activity causes large fluctuations in the mean extension of the filament, especially for very soft filaments with *K*_*s*_*/k*_*m*_ ≪ 1. We also find from additional simulations that motor noise results in uncorrelated spatial domains for *K*_*s*_*/k*_*m*_ = 0.01 while a greater degree of correlation is observed for *K*_*s*_*/k*_*m*_ = 10. Concomitantly, the standard deviation in the mean extension (shown as vertical bars in Figs. 6(a) and 6(b) are more correlated for stiffer filaments than for softer filaments.

### 4.4 Decay of oscillatory strains

To study the decay of localized oscillatory strains in this noisy and active setting, we simulate the model problem analyzed in Section 3.4.5. We impose a low frequency (*ω* ≪ *v*_0_*/σ*), small amplitude (*u*_1_ ≪ *u*_0_(*ℓ*)) oscillation at the free end (*s* = *ℓ*) about the steady stationary extension that is achieved in the absence of this imposed deformation. Thus, the net extension imposed (and measured) at the free end is *u*(*ℓ*)= *u*_0_(*ℓ*)+ *u*_1_ sin(*ωt*). Since the motors do not form permanent cross-links, this steady-state deformation is identical to the deformation attained as the frequency goes to zero. The oscillation at *s* = *ℓ* results in a spatially and temporally varying extension field *u*(*s*) along the contour of the filament. After initial transients have died, this field can be written as the sum of the actively generated steady strain *u*_0_ and an oscillatory strain of amplitude *û*(*s*) ∝ *u*_1_. For small deformations, the ratio *û*(*s*)*/u*_1_ depends only on *s*, the elasticity of the filament, and the motor kinetics. In the frequency-locked limit *û*(*s*)*/u*_1_ oscillates with frequency *ω* - the critical assumption here is that intrinsic oscillatory instabilities are not excited due to motor activity.

Focusing on the amplitude *û* of the oscillatory part of the strain, we compute the spatial variation and fluctuations (averaged over many motor cycles) in the amplitude for each segment as a function of its distance from the free end (*ℓ* − *s*). For sufficiently low frequencies, the results are qualitatively independent of frequency *ω*. For high frequencies, we observed that in some cases localized oscillations were initiated - for these systems the notion of a persistence length breaks down and is not a useful physical measure. Hence, we will focus on the low-frequency results and, in particular, the value *ω* = 10^−3^(*σ/v*_0_).

In the case of an inextensible filament, the oscillatory strain imposed at the free end would propagate an infinitely long distance without any decay in its amplitude. However, for finitely extensible filaments, Equation (30) predicts an exponential decay of strain along the filament. In fact, simulations confirm this feature. Figures 7 (a)-(c) show an exponential decay for the scaled amplitude of the oscillatory extension *û/u*_1_ as a function of distance from the free end for different values of *K*_*s*_*/k*_*m*_ and *p*_on_*/p*_off_. The amplitude *û* does show large fluctuations due to the noise imposed by discrete motor attachment or detachment - something the mean-field theory cannot capture. The length scale over which the elongation decays *λ*_*E*_ is seen to be a strong function of *K*_*s*_*/k*_*m*_, but surprisingly only a weak function of *p*_on_*/p*_off_ in the regime we probed. We hypothesize that this is because the system accessible the dynamical parameter space where the average number of attached motors is small and the contribution to the elastic stiffness due to motor attachment is negligible.

**Fig. 7.**
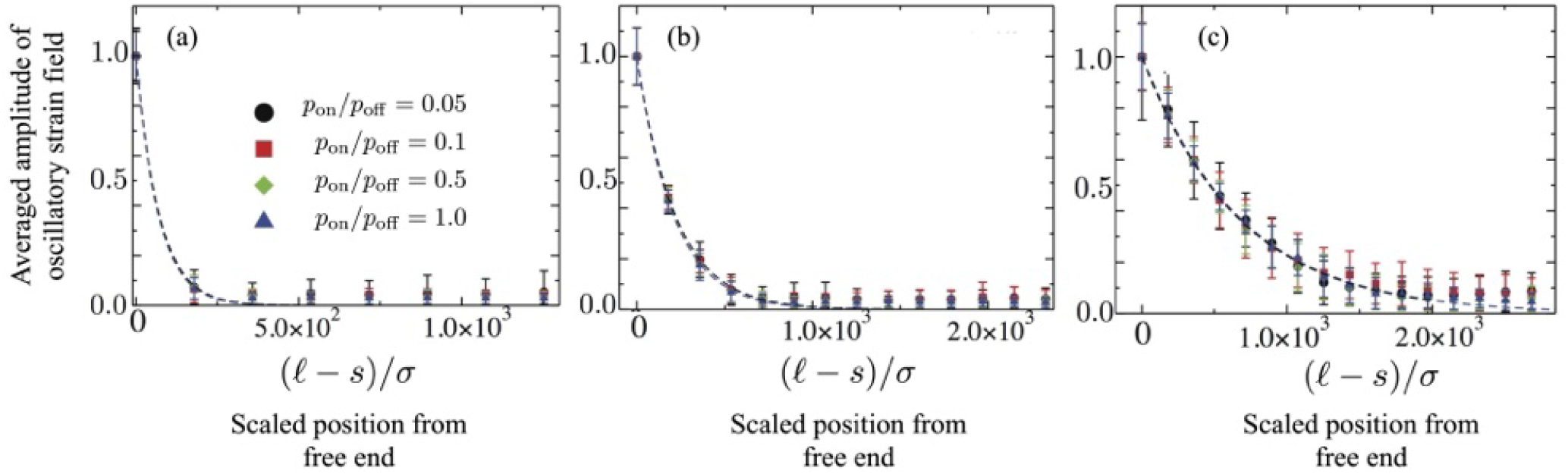
The noise averaged amplitude of spatial oscillatory field of the filaments as a function of position *s*, for different values of the stiffness parameter (a) *K*_*s*_*/k*_*m*_ = 10^−3^, (b) 10^−2^ and (c) 10^−1^. Here *ℓ* is the total length of the composite filament. The dashed curves show exponential fits indicating decay lengths whose value depends on motor kinetics via the ratio of attachment to detachment probabilities as well as on the elasticity contrast. The (dimensionless) forcing frequency is 10^−3^ *v*_0_*/σ*.

### 4.5 Coherence and decay lengths

In summary, we find that the decay of localized, motor-mediated, steady extensions is controlled by *K/k*_*m*_ and the ratio of detachment to attachment probabilities, *p*_off_*/p*_on_, for both stiff (*K*_*s*_*/k*_*m*_ = 10) and soft (*K*_*s*_*/k*_*m*_ = 10^−2^) filaments. The prediction of the mean field is qualitatively consistent with the simulation results for *p*_off_*/p*_on_ ~ 1 with deviations seen for large contrasts between probabilities. Increasing the attachment probability results in a larger fraction of attached motors with sharper gradients near the clamped end and flatter profiles near the free end, consistent with analytical predictions. Motor noise results in large fluctuations in the mean extension, especially for the soft filaments, and also results in uncorrelated spatial domains, as deduced by comparing Figures 8(a) and 8(b).

**Fig. 8.**
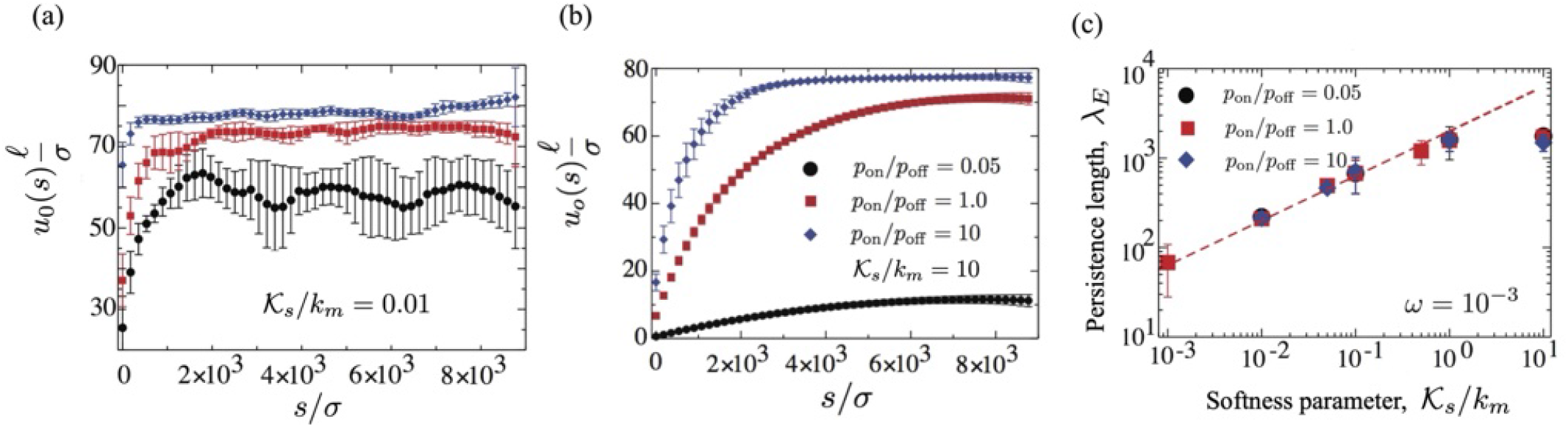
Scaled mean steady extension as a function of position from the clamped end. Vertical bars correspond to the RMS deviation. Extensions are uncorrelated for soft assemblies plotted in (a) for *K*_*s*_*/k*_*m*_ = 0.01 with domains fluctuating independently. In (b) we show results for a stiffer filament with *K*_*s*_*/k*_*m*_ = 10.0 where domains are more correlated. (c) Decay length *λ*_*E*_ as a function of *K*_*s*_*/k*_*m*_ for various *p*_on_*/p*_off_. For small ratios (weak elasticity), simulations agree with the theoretical scaling (dashed line). We see evidence of system-size saturation for large values of *K*_*s*_*/k*_*m*_ as the simulations deviate from the mean-field prediction. The (dimensionless) forcing frequency is 10^−3^*v*_0_*/σ*.

For a fixed *ω*, we estimate the decay length by fitting *û*(*ℓ* − *s*) to an exponential function for a range of *K*_*s*_*/k*_*m*_ and *p*_on_*/p*_off_. Our calculations show an increase in *λ*_*E*_ as *K*_*s*_*/k*_*m*_ varies from 10^−3^ to 10. We plot the computationally calculated *λ*_*E*_ for the noisy system as a function of *K /k*_*m*_ in Fig. 8(c). Our simulation results show excellent agreement with this prediction for almost all of the range explored in *K*_*s*_*/k*_*m*_. For very stiff filaments, we find that the computational value deviates from the dashed line and seems to saturate to a constant, possibly due to finite system-size effects.

The analytical prediction (38) can be simplified for our system without permanent linkers by taking the limit *G*_pas_ = 0. Recasting the resulting expression yields

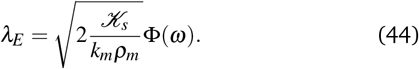

where the function Φ incorporates the dependence on dimensionless frequency. We also calculated the decay length for the imposed frequency (dimensionless) *ω* = 10^−2^ and found qualitatively similar results. For example, when *K*_*s*_*/k*_*m*_ = 1, the decay length is proportionally smaller than for *ω* = 10^−3^ due to the dependence of *χ* on the frequency. For large frequencies, the homeostatic response is not valid and the concept of a decay length loses its significance.

## 5 Concluding remarks

The scale over which active deformations persist in a fluctuating environment is a fundamental question that arises in the study of living, active, and activated matter. Here, this question is addressed in the context of a composite assembly consisting of cross-linked elastic filaments that can be stretched and sheared by ordered active motors. We present a mean-free theory valid in the noiseless weakly soft limit, followed by detailed Brownian multi-particle collision dynamics-based numerical simulations for noisy and moderately soft composites. Both theory and simulations show that even when the filaments comprising the composite are stiff and extensibility may be negligible locally, it cannot be ignored globally. This is because the length scale over which deformations persist is controlled by the competition between filament extensibility and shear stiffening conferred by the permanent passive cross links and temporary attachments formed by the active motors. The mean field theory developed in this work provides analytical expressions for this length scale in the frequency-locked homoeostatic limit. We find that the persistence or the decay length depends on motor kinetics and duty ratio, motor elasticity, and filament extensibility. For fixed motor properties, the persistence length for extensional deformations is controlled by the ratio of the passive elastic modulus of the composite to an effective active motor-generated shear modulus. In our discrete model, the coupling between motor activity and filament dynamics is assumed at the level of individual motors, as in the case of in vitro experiments. This allows us to elucidate the role of complete two-way coupling between motor activity and filament dynamics with minimal assumptions at the macroscopic level.

Surprisingly, the predictions of the mean field theory agree qualitatively with the results from the stochastic discrete filament-motor model, even for moderately strong noise. For strong athermal motor noise, we predict rich dynamical features, including the modification of the motor duty-ratio with filament elasticity, and localized regions of coherent oscillations. Motor noise resulting from discrete binding and unbinding events and due to finite motor numbers results in a fuzzy boundary exhibiting intermittency between these two stable regimes. This suggests that localized, bounded oscillations may arise naturally in long filaments. Taken together, our analytical and computational results also provide insight into how frequency-dependent decay lengths and noise may enable oscillation and synchronization. Consider, for example, Figure 1, and let a local static strain field be generated at location I by an externally imposed force or stretch. As described in our theory, the localized strain at location I will result in a strain field at location II if the distance between them is less than *λ*_*E*_. The propagated strain field could, under certain conditions (such as when boundary constraints are present), trigger local oscillations at location II. In a second scenario, a local oscillatory strain field at location I may emerge naturally due to local instability. Let this emergent field be characterized by frequency *ω*_*I*_ and mean *U*_*I*_. If the separation distance between locations I and II is less than *λ*_*E*_, regions I and II may eventually oscillate synchronously with frequency *ω*_*I*_. A route to synchronization is also suggested by our simulations with motor noise and specifically by Fig. 8(a) for soft filaments. Motor noise may provide significant temporal variations in the strain fields in I and II, and the two regions eventually oscillate with a self-determined mean frequency.

Although our simulations show deviations from noiseless mean field predictions due to the presence of strong discrete noise caused by the variations in motor activity, several qualitative features are surprisingly still retained. Specifically, for fixed motor attachment and detachment rates, the decay length is set by the ratio of the passive elastic modulus of the constituent filament to the active shear modulus generated by the attached motors, even in the presence of noise. This confirms a finite range for correlated activity that might be relevant for various types of ordered active matter, including eukaryotic flagella, reconstituted filament-motor aggregates, and fiber networks ^16,65,66^ particularly in the isostatic limit. We expect that an extension of the analysis proposed here may also be relevant to quantifying persistence lengths in active and biological polymers in both free and constrained geometries ^22,70–72^. A clear and exciting extension would be to extend our analysis to recent work on designing bioinspired and synthetic active cross-linked networks ^73,74^.

## Author contributions statement

LM, AG, and RC designed the research. AG and LM contributed to formulating the theory and analysis. RC formulated the stochastic model and performed simulations. AG and RC analyzed the data. AG, RC and LM wrote the paper.

## Acknowledgments

The Newtonian model analyzed in the ESM is adapted from work by AG on general viscoelastic models supported by grant NSF-MCB-2026782.

## Additional information

### Competing interests

The authors declare no competing interests.

## Notes

### Competing Interest Statement

The authors have declared no competing interest.

### Summary of Updates

Discussion sections have been revised. New equations have been added, and figures modified. Typographical errors have also been corrected in the revised version.

