## Supplementary Information for "Effective elasticity and persistence of strain in active filament-motor assemblies"

### SUPPLEMENTARY INFORMATION
### Effective elasticity and persistence of strain in structured filament-motor assemblies

Arvind Gopinath[a‡], Raghunath Chelakkot[b‡], L Mahadevan[c]

#### 1. Motor driven instabilities of rigid assemblies

##### 1.1 Minimal model for a one-segment system

Figure 1(a) is a discrete version of the weakly extensible continuum filament analyzed in the main manuscript. Specifically, a continuous filament of length $\ell$ is treated as a chain of identical $N \gg 1$ rigid segments, each of length $\ell_s$ such that $\ell \gg \ell_s$. Each rigid segment is connected to its two neighbors by linear elastic springs. Global extension i.e, stretching is accommodated by the extension of each connecting spring. An array of molecular motors (purple) grafted to the underlying surface (black line) attaches stochastically only to the rigid segments and displaces them relative to their rest position. Shear resistance to sliding deformations is provided by passive elastic springs (in blue). These springs crosslink rigid segments to the underlying substrate and can only stretch horizontally. The composite assembly (segments + passive links + motors) is permeated by a Newtonian fluid that provides a medium for viscous dissipation.

To understand the elastic properties of the whole connected filament/chain, we begin by focusing on the dynamics of a single interior segment. We choose a segment (S) with index $k$ located far from the ends of the composite filament. The reference zero-strain – equivalently, zero-extension – rest state for S corresponds to the connecting springs on either side being in the unextended state. A strained state corresponds to the springs deformed from this rest position. We specify that the elastic restoring forces that impede the lateral motion of the $k^{\text{th}}$ segment displacing relative to its neighboring segments (with indices $k-1$ and $k+1$) can be consolidated into a total elastic force from an *effective* linear anchoring spring with stiffness $K_s$ (Figure 1 (b)). Additional elastic resistances to the motion of segment S comes from the passive crosslinks and attached motors. When passive linkers and active motors are both absent (when $\rho_N = 0$ and $\rho_m = 0$), the effective spring constant $K_s \sim K_{\text{pas}}/(\ell_s b)$. When passive linkers are present but active motors are absent ($\rho_N > 0$ and $\rho_m = 0$), $K_s \sim K_s(K_{\text{pas}}, G_{\text{pas}})$. Active motors contribute to $K_s$ as they provide time varying elastic resistance when transiently attached to the filament.

The equations for the dynamics of this one-segment filament-motor assembly (Figure 1(b)) are readily derived using force balances coupled with specifications for motor attachment and detachment. Two scaled variables quantify the state of the (single) filament-motor assembly: (a) the attached motor density fraction $N$, and (b) the mean motor strain $Y$. The evolution of these variables follows

$$\frac{dN}{dt} = (1-N) - \Psi \mathscr{W}(\mathscr{E}, Y) N, \tag{1}$$

$$\frac{dY}{dt} = -\frac{dU}{dt} + \mathscr{V}\Psi(\mathscr{F} - Y) + (\mathscr{D} - Y)\left(\frac{1-N}{N}\right) \tag{2}$$

$$\frac{dU}{dt} = \frac{1}{\mathscr{A}}(-U + \beta N Y) \tag{3}$$

with all variables scaled as explained in the main text. The scaled spring extension $U \equiv u/\delta_m$ with $u$ being the dimensional extension of the linear spring ,and $\delta_m$ being the characteristic size of the motor step. Equations (1)–(3) feature 4 dimensionless positive parameters $\mathscr{V}$, $\mathscr{F}$, $\mathscr{D}$ and $\beta$ that are discussed in detail in the main manuscript. In the main text, we ignore the effects of viscous dissipation due to the drag from the ambient fluid. Here, we retain the effect of medium viscosity and consider motion in an ideal Newtonian fluid. Equation (3) therefore involves a fifth parameter $\mathscr{A} > 0$ that quantifies the effect of external fluid drag and is linear in $\ell_s$ and in the fluid viscosity $\mu$.

[a] *Department of Bioengineering, University of California Merced, Merced, CA, USA.*
[b] *Department of Physics, Indian Institute of Technology Bombay, Mumbai, India.*
[c] *Department of Physics, Department of Organismic and Evolutionary Biology, and School of Engineering and Applied Sciences, Harvard University, Cambridge, MA, USA.*
[‡] *These authors contributed equally to this work*

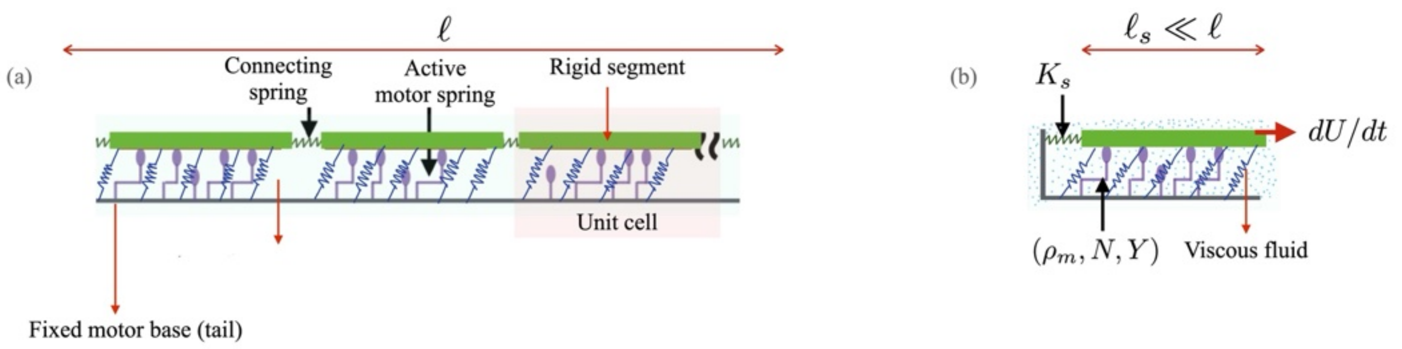


Fig. 1 (a) Schematic of the discrete filament-motor assembly, here modeled as a connected chain of $N$ identical rigid motor-filament segments of length $\ell_s$. Adjacent segments are connected to each other via linear elastic springs that accommodate lateral displacements. Each segment (green) is animated by the action of arrayed molecular motors (illustrated in purple) that may be in one of two states: attached to the segment or detached. Passive linearly elastic cross-linkers that confer shear resistance are shown in blue. (b) The resistance to motion of the test segment from the spring connecting it to neighbors can be combined into an effective elastic resistance with spring constant $K_s$.

The main text addresses the limit $\mathscr{A} = 0$. More complicated forms of fluid drag such as for ambient viscoelastic ambient media have been analyzed in earlier work[1]. When $\mathscr{V} \ll 1$, motor extension is dominated by the pre-strain of *attaching* motors. When $\mathscr{D} \ll 1$, the motor pre-strain has negligible influence.

##### 1.2 Linear stability of the stationary base state in the absence of viscous drag

When $K_s > 0$, the segment cannot translate freely due to the anchoring spring. Under active forcing, the extension $U$ is bounded and therefore either a time-independent constant or a time-periodic function. We first identify the stable static base state by setting the time derivatives in (1)–(3) to zero. This gives us the values for the three variables $(N_0, Y_0, U_0)$ in the base state:

$$(N_0,\ Y_0,\ U_0) = \left( \frac{1}{1+\Psi\mathscr{W}_0},\ \frac{\mathscr{V}\mathscr{F}+\mathscr{D}\mathscr{W}_0}{\mathscr{V}+\mathscr{W}_0},\ \beta N_0 Y_0 \right). \tag{4}$$

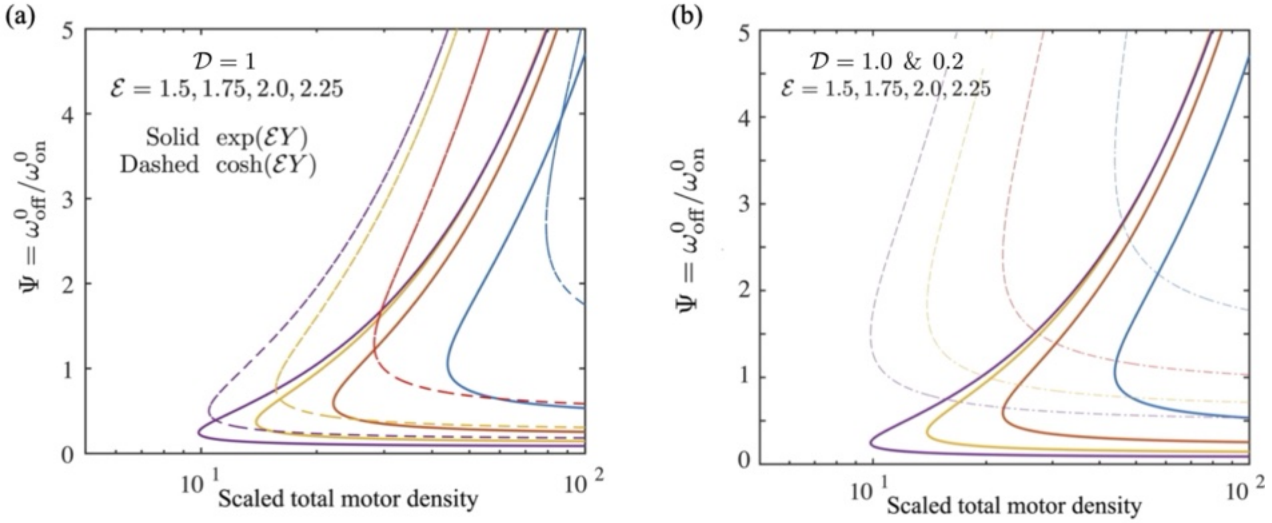


Fig. 2 (a) Neutral stability curves for the case where there is no fluid drag with $\mathscr{A} = 0$, $\mathscr{D} = 1.0$, and $\mathscr{V} = 0$ here. Two forms of the detachment function are shown here, $\mathscr{W} = \exp(\mathscr{E}|Y|)$ (solid curves) and $\mathscr{W} = \cosh(\mathscr{E}Y)$ (dashed curves). The curves correspond (from top to bottom) to $\mathscr{E} = 1.5, 1.75, 2.0$ and $2.25$. We see that for each value of $\mathscr{E}$, there is a critical value of $\Psi(\mathscr{E})$ below which the base state is linearly stable and no oscillations ensue. For a fixed value of $\beta$, oscillations are stable for a range $\Psi_{\rm L}(\beta) < \Psi < \Psi_{\rm U}(\beta)$. (b) Change in the shape of the neutral stability curves as $\mathscr{D}$ changes. Solid curves correspond to $\mathscr{D} = 1.0$, and dashed curves correspond to $\mathscr{D} = 0.2$, and in both cases we have chosen $\mathscr{W} = \exp(\mathscr{E}|Y|)$.

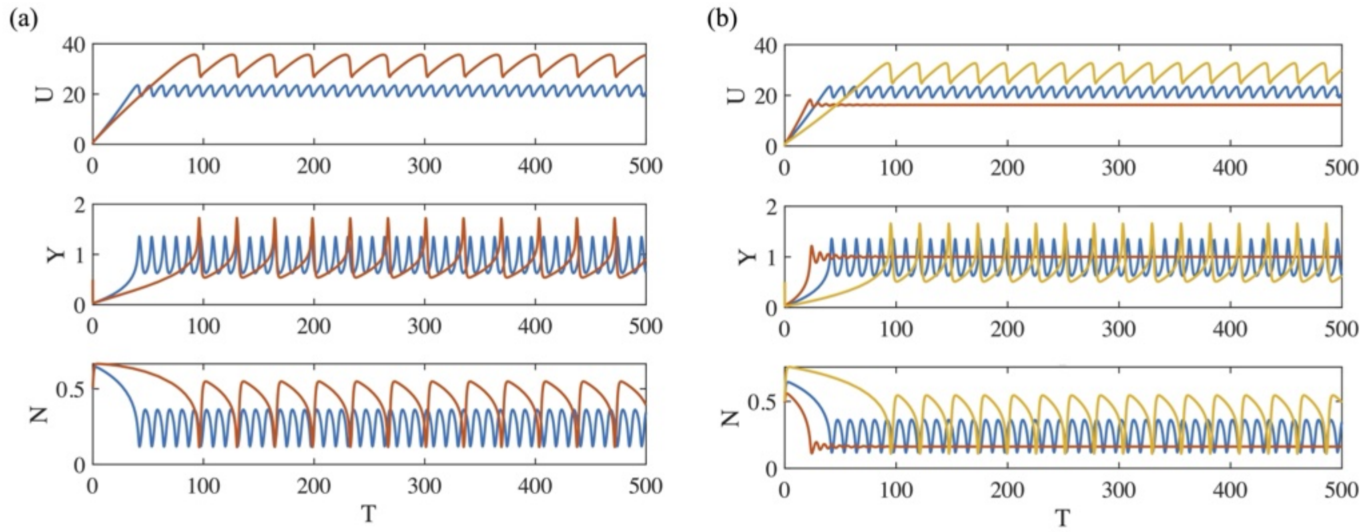


Fig. 3 Stable oscillations in the presence of ambient Newtonian fluid exerting linear fluid drag. The parameters corresponding to these results are $\mathscr{A} = 2.0$, $\mathscr{D} = 1$, $\mathscr{V} = 0$, and $\beta = 100$. (a) Oscillations in time $T$ as exhibited by $U(T)$, $N(T)$ and $Y(T)$. The blue curves correspond to $\mathscr{W} = \exp(\mathscr{E}|Y|)$, and red curves correspond to $\mathscr{W} = \cosh(\mathscr{E}Y)$. We set $\Psi = 0.5$ and $\mathscr{E} = 2$. (b) Oscillations may be modulated and even suppressed by increasing $\Psi$. This trend is demonstrated here by varying $\Psi$ and examining the solutions. The curves here correspond to $\Psi = 0.3$ (yellow), $\Psi = 0.5$ (blue), and $\Psi = 0.8$ (red). The detachment function $\mathscr{W} = \exp(\mathscr{E}|Y|)$ with $\mathscr{E} = 2$. Note that when $\Psi$ is increased to $0.8$, the system does not oscillate and instead stabilizes to a steady stationary state.

To study the stability of this base state to infinitesimal perturbations, we linearize equations (1)-(3) about $(N_0, Y_0, U_0)$, and examine the eigenvalue structure of the resulting coupled ODE system. For $\mathscr{A} = 0$, (the focus of the main manuscript), linear stability analysis indicates that the base state may be linearly unstable to small perturbations that drive the system to a stable periodic oscillatory state via a supercritical Hopf-Poincare-Andronov bifurcation. To further quantify the emergent oscillatory states and the parameter range over which the instability manifests, we focus on the limit where the third term on the right hand side of equation (2) dominates the second term on the right hand side. This limit has also been studied in previous work[2]. Setting $\mathscr{D} = 0$ and $\mathscr{V} \neq 0$, and completing the linear stability analysis, we obtain expressions for the critical frequency at onset $\omega_c$

$$\omega_c = \omega_{\text{on}}^0 \sqrt{\left( \frac{(\Psi\mathscr{V} + \Psi\mathscr{W}_0)(1 + \Psi\mathscr{W}_0) + Y_0 \Psi \mathscr{W}_0' N_0^{-1}}{1 + \beta N_0} \right)}, \tag{5}$$

and the equation for the neutral stability curve

$$\beta Y_0 \Psi \mathscr{W}_0' N_0 = \Psi\mathscr{V} + (1 + 2\Psi\mathscr{W}_0) + \beta N_0 (1 + \Psi\mathscr{W}_0). \tag{6}$$

The biophysical origin of these oscillations is evident upon examination of the eigenvectors and eigenvalues of the linearized equations. Oscillations result from a positive feedback process due to the emergence of *negative effective* spring constants and/or negative *active* friction coefficients. Active friction arises from dissipation that occurs when the motors detach and relax to their unbound state. For large values of $\Psi$, the number of motors attached may be insufficient to drive the oscillations. In contrast, for small $\Psi$, too many attached motors increase the net elastic active resistance and associated high (active) dissipation/friction coefficients. In the stable oscillating state between these limits, the active energy input by the motor attachment balances the dissipation due to motor friction.

When motors are adiabatically coupled to the dynamics of the segment, that is, when $\dot{Y} \approx \Psi Y \mathscr{W}$, the critical frequency and effective motor friction are, respectively,

$$\omega_c \approx \sqrt{\frac{K_s(\omega_{\text{on}}^0 + \omega_{\text{off}}^0 \mathscr{W}_0)}{\rho_m k_m N_0 \delta_m (dY/d\dot{U})_{\dot{U}=0}}}, \quad \text{and} \tag{7}$$

$$\eta_{\text{active}} \sim \rho_m k_m \delta_m N_0 (dY/d\dot{U})_{\dot{U}=0}. \tag{8}$$

In equations (7) and (8) the dot implies the time derivative.

**Case 2:** In the complementary limit $\mathscr{V}=0$ and $\mathscr{D}\neq 0$, stability analysis likewise provides the equation for the neutral stability curve separating the linear stable and unstable regions of the phase-space,

$$0=(1+2\Psi\mathscr{W}_0)-\beta Y_0\Psi N_0\mathscr{W}_0'+\beta N_0(1+\Psi\mathscr{W}_0). \tag{9}$$

Equation (9) can be used to estimate the critical value of $\beta$ (or equivalently, the critical density of motors $\rho_m=\rho_m^c$ that ensures linear instability. The frequency at criticality is

$$\omega_c=\omega_{\text{on}}^0\sqrt{\frac{\Psi\mathscr{W}_0(1+\Psi\mathscr{W}_0)}{(1+G_{\text{act}}N_0/K_s)}}. \tag{10}$$

Equations (5)-(10) highlight the result that *the motor filament assembly can spontaneously oscillate even without viscosity and dissipation due to drag from an external fluid medium*.

##### 1.3 Linear stability of the stationary base state in the presence of fluid drag

The effects of Newtonian (linear, low Reynolds number) and non-Newtonian fluid drag have been extensively analyzed in previous work[1]. Here we provide a summary of pertinent results for the purely viscous low-Reynolds number case. We consider a model system with $\mathscr{V}=0$ so that pre-strain dominates. The equations reduce to the form

$$\frac{dN}{dt}=(1-N)-\Psi\mathscr{W}(\mathscr{E},Y)N, \quad \frac{dY}{dt}=-\frac{dU}{dt}+(\mathscr{D}-Y)\left(\frac{1-N}{N}\right), \quad \text{and} \quad \mathscr{A}\frac{dU}{dt}+U=\beta NY. \tag{11}$$

Linear stability analysis of (11) shows that stable oscillatory states can emerge at critical points via a Hopf-Andronov-Poincare bifurcation for high motor density, regardless of the value of $\mathscr{A}$. The associated frequencies at onset and locus of critical points satisfy

$$\mathscr{A}\left(\frac{\omega_c}{\omega_{\text{on}}^0}\right)^2 = [(1+2\Psi\mathscr{W}_0)+\mathscr{A}\Psi\mathscr{W}_0(1+\Psi\mathscr{W}_0)-\beta Y_0\Psi N_0\mathscr{W}_0'+\beta N_0(1+\Psi\mathscr{W}_0)], \quad \text{and} \tag{12}$$

$$\omega_c = \omega_{\text{on}}^0\sqrt{\frac{\Psi\mathscr{W}_0(1+\Psi\mathscr{W}_0)}{(1+\beta N_0)+\mathscr{A}(1+2\Psi\mathscr{W}_0)}}. \tag{13}$$

These equations simplify further when motors are adiabatically coupled to the motion of the segment, that is, when $dY/dt=\Psi Y\mathscr{F}(\mathscr{E},Y)$. In this case, we find

$$\beta N_0Y_0\Psi\mathscr{W}_0' = (\beta N_0+\mathscr{A}(2\Psi\mathscr{W}_0+\Psi Y_0\mathscr{W}_0')(1+\Psi\mathscr{W}_0)+(2\Psi\mathscr{W}_0+\Psi Y_0\mathscr{W}_0') \tag{14}$$

$$\omega_c = \omega_{\text{on}}^0\sqrt{\frac{(2\Psi\mathscr{W}_0+\Psi Y_0\mathscr{W}_0')(1+\Psi\mathscr{W}_0)}{\beta N_0+\mathscr{A}(2\Psi\mathscr{W}_0+\Psi Y_0\mathscr{W}_0')}}. \tag{15}$$

As expected, since $\mathscr{A}>0$, fluid drag decreases the frequencies of emergent oscillatory states at onset (for parameters close to the critical parameters) relative to the case without drag. The frequencies of oscillatory states when parameters are far from onset are studied in more detail in our previous work[1].

#### 2. Mean-field model

##### 2.1 Decay length depends on the compliance function $\chi$

We start with equations (33)-(35) for $(\hat{U},\hat{Y},\hat{N})$ in the main text. When the forcing at $s=\ell$ is periodic with time dependence $\Omega(\omega,t)=\exp(i\omega t)$, the linearized equations become

$$\begin{aligned}\frac{\partial^2 \hat{U}}{\partial \bar{s}^2}-\hat{U}(1+\beta\chi) &= 0 \quad (16)\\ \hat{N}(i\omega+1+\Psi\mathscr{W}_o)+\Psi N_o\mathscr{W}_o'\hat{Y} &= 0 \quad (17)\\ i\omega\hat{U}+\hat{Y}(i\omega+\mathscr{V}\Psi+\Psi\mathscr{W}_o)+\hat{N}\left(\frac{\mathscr{D}-Y_0}{N_0^2}\right) &= 0 \quad (18)\end{aligned}$$

where the linear (complex) viscoelastic response function $\chi=-(N_0\hat{Y}+Y_0\hat{N})/\hat{U}$ is determined by the motor kinetics model. Equation (16) is a second-order linear ODE for the spatially dependent complex valued function $\hat{U}(s)$ with constant but complex coefficients. The equation is homogeneous since the term independent of $\hat{U}$ is identically zero. It is instructive to go back to the dimensional form of Equation (16), that is,

$$\ell_E^2\frac{\partial^2\hat{U}}{\partial s^2}-\hat{U}(1+\beta\chi)=0. \quad (19)$$

Setting $\hat{U}=A+iB$, and separating the compliance function into its real and imaginary components $\chi=\mathscr{R}+i\mathscr{I}$, we obtain the following coupled second-order linear ODE system with real coefficients and real boundary conditions:

$$\begin{aligned}\ell_E^2A''-A(1+\beta\mathscr{R})+B\beta\mathscr{I} &= 0, \text{ and} \quad (20)\\ \ell_E^2B''-A\beta\mathscr{I}-B(1+\beta\mathscr{R}) &= 0. \quad (21)\end{aligned}$$

In equations (20) and (21) primes denote differentiation with respect to $s$. Next, we define a column vector, $\mathbf{X}=(A\ B)^T$ and write Equations (20) and (21) as a matrix product $\mathbf{QX}=\mathbf{0}$. Setting $\mathbf{X}=(A^*\ B^*)^T\exp(-qs)$ (where $q$ is complex), we find

$$\begin{aligned}0 &= A^*(1+\beta\mathscr{R}-\ell_E^2q^2)-B^*\beta\mathscr{I}, \quad \text{or} \quad \frac{A^*}{B^*}=\frac{\beta\mathscr{I}}{(1+\beta\mathscr{R}-\ell_E^2q^2)}, \quad \text{and} \quad (22)\\ 0 &= A^*\beta\mathscr{I}+B^*(1+\beta\mathscr{R}-\ell_E^2q^2), \quad \text{or} \quad \frac{A^*}{B^*}=-\frac{(1+\beta\mathscr{R}-\ell_E^2q^2)}{\beta\mathscr{I}} \quad (23)\end{aligned}$$

which when combined provides the equation for $\lambda=(\ell_Eq)^2$,

$$\lambda^2-2\lambda(1+\beta\mathscr{R})+(\beta^2\mathscr{I}^2+(1+\beta\mathscr{R})^2)=0. \quad (24)$$

The solutions to (24) are the set of all values of $\lambda$, denoted here by $\lambda_\pm$, and satisfy

$$\begin{aligned}\lambda_+=\ell_E^2q_+^2 &= (1+\beta\mathscr{R})+i\beta\mathscr{I}=(1+\beta\chi), \quad \text{and} \quad q_+=\ell_E^{-1}\sqrt{(1+\beta\mathscr{R})+i\beta\mathscr{I}}=\ell_E^{-1}\sqrt{z}=\ell_E^{-1}\sqrt{x+iy} \quad (25)\\ \lambda_-=\ell_E^2q_-^2 &= (1+\beta\mathscr{R})-i\beta\mathscr{I}=(1+\beta\bar{\chi}), \quad \text{and} \quad q_-=\ell_E^{-1}\sqrt{1+\beta\bar{\chi}}=\ell_E^{-1}\sqrt{\bar{z}}=\ell_E^{-1}\sqrt{x-iy}. \quad (26)\end{aligned}$$

Note that $\bar{\chi}$ is the complex conjugate of $\chi$, $x=1+\beta\mathscr{R}$, and $y=\beta\mathscr{I}$. Focusing on equation (25), we find that the principal square root of $z$ is given by

$$\begin{aligned}\sqrt{z} &= (\alpha_1+i\alpha_2) \quad \text{where} \quad (27)\\ \alpha_1 &= \sqrt{\frac{\sqrt{(1+\beta\mathscr{R})^2+(\beta\mathscr{I})^2}+(1+\beta\mathscr{R})}{2}} \quad \text{and} \quad \alpha_2=\frac{\mathscr{I}}{|\mathscr{I}|}\sqrt{\frac{\sqrt{(1+\beta\mathscr{R})^2+(\beta\mathscr{I})^2}-(1+\beta\mathscr{R})}{2}}. \quad (28)\end{aligned}$$

Therefore the two relevant values of $q_+$ are $q_1 = +\ell_E^{-1}(\alpha_1 + i\alpha_2)$ and $q_2 = -\ell_E^{-1}(\alpha_1 + i\alpha_2)$. Using the expansion

$$\exp(-q_1 s) = \exp\left(-s/\ell_E \alpha_1^{-1}\right)\left(\cos(\ell_E^{-1}\alpha_2 s) - i\sin(\ell_E^{-1}\alpha_2 s)\right) \tag{29}$$

we identify the decay length

$$\lambda_E = \ell_E \alpha_1^{-1}. \tag{30}$$

Similarly noting that $\bar{z}$ and $z$ differ in just the sign of the negative part, we identify the two solutions to $q_-$, $q_3 = +\ell_E^{-1}(\alpha_1 - i\alpha_2)$ and $q_4 = -\ell_E^{-1}(\alpha_1 - i\alpha_2)$.

The complete set of $q$ values that solve (24) are $(q_1, q_2, q_3, q_4)$. The solution to $\hat{U}$ is a combination of independent linear modes corresponding to these values of $q$, and the appropriate coefficients (vectors) are determined by the boundary conditions at the two ends ($s = 0$ and $s = \ell$). The presence of $\cos(\ell_E^{-1}\alpha_2 s)$ and $\cos(\ell_E^{-1}\alpha_2 s)$ terms confirms that the fields are oscillatory with the amplitude decaying over the length scale $\lambda_E$. The wavelength of the oscillations is determined to be $\ell_E \alpha_2^{-1}$. Both these length-scales depend on motor kinetics and on the imposed frequency.

##### 2.2 Determination of the linear compliance function, $\chi$

The compliance function $\chi = -(N_0\hat{Y} + Y_0\hat{N})/\hat{U}$, in Eqn. (19), is dependent on motor kinetics and properties. Note that the frequency $\omega$ is dimensionless in (16)-(18). Here we show how $\chi$ is evaluated in two limits.

###### 2.2.1 Expression in the limit $\mathscr{V} = 0$

In this limit, we have the following:

$$(Y_0, N_0) = (\mathscr{D}, (1 + \Psi\mathscr{W}_0)^{-1}) \tag{31}$$

Equations (17) and (18) reduce to

$$\hat{N}(i\omega + 1 + \Psi\mathscr{W}_o) + \Psi N_o \mathscr{W}_o' \hat{Y} = 0 \tag{32}$$

$$i\omega\hat{U} + \hat{Y}(i\omega + \Psi\mathscr{W}_o) + \hat{N}\left(\frac{\mathscr{D} - Y_0}{N_0^2}\right) = 0 \tag{33}$$

which we combine to write

$$\chi(\omega) = -(N_0\hat{Y} + Y_0\hat{N})/\hat{U} = -\mathscr{D}\left(\frac{\Psi N_0 \mathscr{W}_0'}{1 + i\omega + \Psi\mathscr{W}_0}\right)\left(\frac{i\omega}{i\omega + \Psi\mathscr{W}_0}\right) + \left(\frac{1}{1 + \Psi\mathscr{W}_0}\right)\left(\frac{i\omega}{i\omega + \Psi\mathscr{W}_0}\right). \tag{34}$$

Separating $\chi = \mathscr{R}(\omega) + i\mathscr{I}(\omega)$ into its real and imaginary components, we find

$$\mathscr{R}(\omega) = +\left[\frac{\omega^2}{(1 + \Psi\mathscr{W}_0)(\Psi^2\mathscr{W}_0^2 + \omega^2)}\right] - \mathscr{D}\left[\frac{\omega^2\Psi\mathscr{W}_0'(1 + 2\Psi\mathscr{W}_0)}{(\Psi^2\mathscr{W}_0^2 + \omega^2)((1 + \Psi\mathscr{W}_0)^2 + \omega^2)(1 + \Psi\mathscr{W}_0)}\right] \tag{35}$$

$$\mathscr{I}(\omega) = +\left[\frac{\omega\Psi\mathscr{W}_0}{(1 + \Psi\mathscr{W}_0)(\Psi^2\mathscr{W}_0^2 + \omega^2)}\right] - \mathscr{D}\left[\frac{\omega\Psi\mathscr{W}_0'(\Psi\mathscr{W}_0 + \Psi^2\mathscr{W}_0^2 - \omega^2)}{(\Psi^2\mathscr{W}_0^2 + \omega^2)((1 + \Psi\mathscr{W}_0)^2 + \omega^2)(1 + \Psi\mathscr{W}_0)}\right] \tag{36}$$

Noting that $\Psi\mathscr{W}_0 = (\omega_{\mathrm{off}}^0/\omega_{\mathrm{on}}^0)\mathscr{W}_0$, we see that the duty ratio in the deformed state is related to the (base-state) duty ratio $\tau$ through

$$\tau \equiv \frac{1}{1 + \Psi\mathscr{W}_0} = \frac{\tau_a}{\tau_a + \tau_d}. \tag{37}$$

Here $\tau_a$ and $\tau_d$ are the ensemble-averaged and load-dependent attached and detached times over a mechanochemical cycle of a motor evaluated at the base state.

For fixed imposed frequency (sufficiently small to maintain homoeostatic conditions), $\tau$ and $\mathscr{D}$ together determine the complex viscoelasticity of the coupled motor aggregates. The condition $\omega \ll \Psi\mathscr{W}_0$ implies $\omega \ll (1 + \Psi\mathscr{W}_0)$, and is therefore the limit $\tau^{-1} \gg \omega$. Since $\Psi\mathscr{W}_0 > 0$ always, $\omega \gg (1 + 2\Psi\mathscr{W}_0)$ implies $\omega \gg (1 + \Psi\mathscr{W}_0)$, and is therefore provides the limit

$\tau^{-1} \ll \omega$. Expanding equations (35) and (36) in these two limits we obtain to leading order

$$\mathscr{R}(\omega \ll \tau^{-1}) \sim \frac{\omega^2}{(1+\Psi\mathscr{W}_0)(\Psi^2\mathscr{W}_0^2)}\left(1-\mathscr{D}\frac{\Psi\mathscr{W}_0'(1+2\Psi\mathscr{W}_0)}{(1+\Psi\mathscr{W}_0)^2}\right) \tag{38}$$

$$\mathscr{I}(\omega \ll \tau^{-1}) \sim \frac{\omega}{(1+\Psi\mathscr{W}_0)(\Psi\mathscr{W}_0)}\left(1-\mathscr{D}\frac{\Psi\mathscr{W}_0'}{(1+\Psi\mathscr{W}_0)}\right) \tag{39}$$

$$\mathscr{R}(\omega \gg \tau^{-1}) \sim \left(\frac{1}{1+\Psi\mathscr{W}_0}\right)-\frac{\mathscr{D}}{\omega^2}\left(\frac{\Psi\mathscr{W}_0'(1+2\Psi\mathscr{W}_0)}{1+\Psi\mathscr{W}_0}\right) \tag{40}$$

$$\mathscr{I}(\omega \gg \tau^{-1}) \sim \frac{1}{\omega}\left(\frac{\Psi\mathscr{W}_0}{1+\Psi\mathscr{W}_0}+\mathscr{D}\frac{\Psi\mathscr{W}_0'}{1+\Psi\mathscr{W}_0}\right)-\frac{\mathscr{D}}{\omega^3}\left(\frac{\Psi\mathscr{W}_0'(\Psi\mathscr{W}_0+\Psi^2\mathscr{W}_0^2)}{1+\Psi\mathscr{W}_0}\right) \tag{41}$$

**2.2.2 Expression in the limit $\mathscr{D} = 0$**

We start by recalling the expressions $\mathscr{V} \equiv v_0 k_m(\omega_{\text{off}}^0 F_s)^{-1}$ and $\mathscr{F} \equiv F_s(k_m\delta_m)^{-1}$. When $\mathscr{D} = 0$ we have the base state given by

$$(Y_0, N_0) = \left(\frac{\mathscr{V}\mathscr{F}}{\mathscr{V}+\mathscr{W}_0}, \frac{1}{1+\Psi\mathscr{W}_0}\right) \tag{42}$$

Using (37), we obtain the susceptibility function $\chi(\omega) = -N_0\left(\hat{Y}/\hat{U}\right) - Y_0\left(\hat{N}/\hat{U}\right)$ in terms of motor kinetics parameters

$$\chi(\omega) = i\omega N_0\left[(i\omega+\mathscr{V}\Psi+\Psi\mathscr{W}_0)+\left(\frac{Y_0}{N_0^2}\right)\left(\frac{\Psi N_0\mathscr{W}_0'}{1+i\omega+\Psi\mathscr{W}_0}\right)\right]^{-1}$$

$$-i\omega Y_0\left(\frac{\Psi N_o\mathscr{W}_0'}{i\omega+1+\Psi\mathscr{W}_0}\right)\left[(i\omega+\mathscr{V}\Psi+\Psi\mathscr{W}_0)+\left(\frac{Y_0}{N_0^2}\right)\left(\frac{\Psi N_0\mathscr{W}_0'}{1+i\omega+\Psi\mathscr{W}_0}\right)\right]^{-1}. \tag{43}$$

#### 3. Steady extension of the aggregate in the absence of passive linkers

In the absence of permanent elastic linkers as in the simulation, $\rho_N = 0$ and therefore the passive shear modulus $G_{\text{pas}} = 0$. The governing equations in dimensional form are

$$K_{\text{pas}}\partial_{ss}U_0 + G_{\text{act}}\delta_m N_0 Y_0 = 0, \tag{44}$$

$$N_0 = (1+\Psi\mathscr{F}_0)^{-1}, \tag{45}$$

$$Y_0 = (\mathscr{V}\mathscr{F}+\mathscr{D}\mathscr{W}_0)(\mathscr{V}+\mathscr{W}_0)^{-1} \tag{46}$$

from which we deduce that the steady extension field $U_0(s)$ is quadratic in $s$ and thus *there is no exponential decay length*. When pre-extension of attaching motors, $d_m = 0$ so that $\mathscr{D} = 0$, the solution is

$$U_0(s) = \left(\frac{\rho_m k_m}{K_{\text{pas}}}\right)\left(\frac{\mathscr{V}\mathscr{F}}{\mathscr{V}+\mathscr{W}_0}\right)\left(\frac{\delta_m}{1+\Psi\mathscr{W}_0}\right)s(2\ell - s). \tag{47}$$

The spatial variation in the strain and the extension at the free end are, respectively, given by

$$\frac{U_0(s)}{U_0(\ell)} = \frac{s(2\ell-s)}{\ell^2}, \quad U_0(\ell) = \left(\frac{\rho_m k_m}{K_{\text{pas}}}\right)\left(\frac{\mathscr{V}\mathscr{F}}{\mathscr{V}+\mathscr{W}_0}\right)\left(\frac{\delta_m}{1+\Psi\mathscr{W}_0}\right)\ell^2. \tag{48}$$

#### 4. Estimates for motor kinetics parameters and persistence lengths

In this section, we summarize typical values and experimentally observed ranges for kinetic parameters and mechanical properties of dynein and myosin motors. These values are obtained from studies[3–8] and references therein.

Cytoplasmic dynein is found to exhibit attachment rates of $\sim 5$ 1/s. The detachment rates are force-dependent and range from $0.1 - 1.7$ 1/s. There is evidence that dynein has a slip-bond response under forward loads but ideal force-independent unbinding at backward forces greater than $\sim 2$ pN. Some studies also identify a catch-bond-like response at backward loads. Experimentally obtained values for stall forces are $\sim 1 - 2$ pN for isolated mammalian dynein, increasing to $\sim 4-7$ pN under certain conditions. Motor step sizes are typically $\sim 8$ nm, but with high variability $\sim 4-32$ nm. The duty ratio, that is, the fraction of attached time, varies significantly $\sim 10-20$ %. Axonemal outer arm and inner arm dyneins have been characterized primarily by gliding assays, microtubule sliding, and optical trapping of intact or semi-demembranated axonemes. Individual outer-arm motors (OAD) exhibit stall forces near $\sim 4.7-5$ pN and low duty ratios $\sim 7-8$ %. Inner arm dyneins (IAD) are observed to have a higher duty ratio[8]. Estimates for free velocities (at zero load) for the outer axonemal dyneins are $\sim 1.2$ $\mu$m/s (in *Tetrahymena*, sea urchin) and $\sim 12-1$ $\mu$m/s for inner arm dyneins. In general, there is significant variation in the estimates of the stall forces ranging from $\sim 0.8$ pN to $\sim 9$ pN depending on the study, the type of dynein, and the organism involved. Studies on OAD in sea urchin sperm suggest the following values: single molecule stall forces $\sim 4.7$ pN, zero-load speed $\sim 5$ $\mu$m/s, and step size $\sim 8$ nm. Studies on OAD in *Chlamydomonas* suggest single motor stall force $\sim 5$ pN, maximum gliding velocity $\sim 5$ $\mu$m/s, and step size $\sim 8$ nm at low ATP. For myosin motors, studies on Myosin IIA suggest attachment rates $\sim 10-50$ 1/s and detachment rates $\sim 20-50$ 1/s. For Myosin IIB, estimated attachment rates are $5-20$ 1/s and detachment rates are $\sim 1-5$ (1/s). Stall forces are $\sim 5-6$ pN per motor head for skeletal muscle, and $\sim 2-3$ pN for non-muscle isoforms. Skeletal myosin exhibits a biphasic displacement of $\sim 5-7$ nm, while other isoforms suggest step sizes of $\sim 5-10$ steps. The force-extension is roughly linear (Hookean) over physiological force ranges.

#### Notes and references


1 J. Tamayo, A. Mishra and A. Gopinath (2022) Front. Phys. 10:895536.

2 A.Vilfan and E. Frey (2005) J. Phys. Condens. Matter, 94, 108104.

3 D. D. Hackney (1996) Ann. Rev. Physiology, 58, 731.

4 A. J. Roberts, T. Kon, P. J. Knight, K. Sutoh and S. A (2013) Nat. Rev. Mol. Cell Biol., 14 (11), 713.

5 X. Liu, L. Rao, A. Gennerich (2023) Methods in Molecular Biology, 2623, 221.

6 M. Linari, G. Piazzesi, I. Pertici, J. A. Dantzig, Y. E. Goldman, and V. Lombardi (2020) Biophys. J. 118 (5), 994.

7 E. R. Lecarpentier, *et. al.* (2011) 111 (4), 1096.

8 T. Fujiwara, C. Shingyoji and H. Higuchi (2023) Sci Rep 13, 10514.
